# AP-2α/AP-2β transcription factors are key regulators of epidermal homeostasis

**DOI:** 10.1101/2023.12.03.569763

**Authors:** Hui Zhang, Jackelyn Raymundo, Kathleen E. Daly, Wenjuan Zhu, Bill Senapati, Alexander G. Marneros

## Abstract

AP-2 transcription factors regulate ectodermal development but their roles for epidermal homeostasis in the adult skin are unknown. We find that AP-2α is the predominant AP-2 family member in adult epidermis, followed by AP-2β. Through inactivation of AP-2α, AP-2β, or both in keratinocytes we assessed the effects of a gradient of epidermal AP-2 activity on skin function. We find that (1) loss of AP-2β in keratinocytes is compensated for by AP-2α, (2) loss of AP-2α impairs terminal keratinocyte differentiation and hair morphogenesis, and (3) the combined loss of AP-2α/AP-2β results in more severe skin and hair abnormalities. Keratinocyte differentiation defects precede a progressive neutrophilic skin inflammation. Inducible inactivation of AP-2α/AP-2β in the adult phenocopies these manifestations. Transcriptomic analyses of epidermis lacking AP-2α or AP-2α/AP-2β in keratinocytes demonstrate a terminal keratinocyte differentiation defect with upregulation of alarmin keratins and of several immune pathway regulators. Moreover, our analyses suggest a key role of loss of AP-2α-dependent gene expression of CXCL14 and KRT15 as an early pathogenic event towards the manifestation of skin inflammation. Thus, AP-2α/AP-2β are critical regulators of epidermal homeostasis in the adult skin.

## INTRODUCTION

The AP-2 transcription factor family consists of AP-2α, AP-2β, AP-2γ, AP-2δ, and AP-2ε (Eckert et al., 2005). Mammalian AP-2 proteins (except AP-2δ) have very similar DNA sequence preferences and bind as hetero- or homodimers to the consensus motif GCCNNNGGC (Williams and Tjian, 1991a, b). AP-2γ has been reported to have an important role in surface ectoderm induction and primes p63-dependent keratinocyte maturation during development (Li et al., 2019). However, keratinocyte-specific inactivation of AP-2γ at E14.5 did not result in a skin phenotype (Wang et al., 2008). Functional redundancy between AP-2α and AP-2γ has been reported in the epidermis as well, where inactivation of AP-2α and AP-2γ at ∼E14.5 (using K14Cre mice) resulted in perturbed terminal differentiation of keratinocytes and perinatal lethality (pups died within 24 hours of birth) (Wang et al., 2008). Conditional inactivation of both AP-2α and AP-2β in the early surface ectoderm at ∼E8.5 (using the ectoderm-expressed Cre recombinase transgene Crect) resulted in defects of the craniofacial complex, including clefts in both the upper face and mandible (Van Otterloo et al., 2022). These craniofacial defects caused a perinatal lethality of these conditional mutant mice, which prevented the assessment of the roles of AP-2α and AP-2β for the adult epidermis.

Conditional inactivation of only AP-2α in keratinocytes at ∼E14.5, using K14Cre mice, resulted in their hyperproliferation and impaired terminal differentiation *in vivo* (Wang et al., 2006). This epidermal hyperproliferation was already present at P3, and no skin inflammation was observed at that time (Wang et al., 2006). Moreover, primary keratinocytes lacking AP-2α showed an increased cell proliferation rate *in vitro*, demonstrating that lack of AP-2α leads to hyperproliferation as a primary event that is not a secondary consequence of an inflammatory process (Wang et al., 2006). This study suggested that AP-2α functions to repress epidermal growth factor receptor (*Egfr*) expression in the epidermis and *via* this mechanism promotes the transition from epidermal proliferation to differentiation (Wang et al., 2006).

Notably, the role of AP-2β in the adult skin has not been assessed and it is not known whether AP-2α and AP-2β have distinct or overlapping compensatory functions in the epidermis. In other epithelial tissues, AP-2α and AP-2β can have distinct non-overlapping expression patterns and functions. For example, we found that in the distal nephron of the kidney, AP-2α and AP-2β show a largely non-overlapping expression pattern and function, where AP-2β is critical for the differentiation and function of the distal convoluted tubule and AP-2α regulates medullary collecting duct function (Lamontagne et al., 2022).

Here, we show that AP-2α is the most abundant AP-2 family member in the epidermis, followed by AP-2β and AP-2γ. By generating mice that lack AP-2β, AP-2α, or both selectively in keratinocytes, we compared distinct mouse mutants with a progressive overall reduction of AP-2 activity in the epidermis and found increasingly more severe primary keratinocyte differentiation defects. We show that the keratinocyte differentiation defects, coupled with hyperproliferation and reduced expression of cell adhesion genes, are associated with a subsequent progressive neutrophilic skin inflammation with upregulated gene expression of Th17 cytokine, complement, and inflammasome genes. Notably, mice lacking both AP-2α and AP-2β in keratinocytes had a more severe differentiation defect than mice lacking only AP-2α, and these double mutant mice also developed a more severe skin inflammation with earlier onset than mice lacking only AP-2α. Similarly, inducible inactivation of AP-2β, AP-2α, or AP-2α/AP-2β in the adult recapitulated these findings. Thus, induction of epidermal hyperproliferation and impairing terminal differentiation by targeting AP-2 activity in keratinocytes is associated with a progressive subsequent neutrophilic skin inflammation. The epidermal gene expression changes in the inflammed skin of mice lacking AP-2α/AP-2β showed a partial overlap with the expression changes seen in keratinocytes of patients with psoriasis or atopic dermatitis. Analyses of differentially expressed genes from (1) mice lacking AP-2α/AP-2β in early surface ectoderm (at E10.5), (2) AP-2α-dependent human keratinocyte genes, (3) transcriptomes from epidermis of our *K14Cre^+^Tfap2a^fl/fl^Tfap2b^fl/fl^* mice with skin inflammation, and (4) human scRNA-seq data from keratinocytes derived from psoriasis and atopic dermatitis lesional skin implicate loss of AP-2α-dependent gene expression of CXCL14 and KRT15 in keratinocytes as early pathogenic events that precede the manifestation of skin inflammation.

Overall, our findings establish AP-2α/AP-2β as critical regulators of epidermal homeostasis and keratinocyte differentiation processes in the adult skin. Moreover, they suggest that therapeutic approaches to normalize keratinocyte proliferation and differentiation may inhibit skin inflammation.

## RESULTS

### AP-2α is the most abundant AP-2 family member in the epidermis

Analysis of scRNA-seq data shows that AP-2α and AP-2β are expressed in the developing epidermis at E12.5 (Figure S1A) and that their expression is maintained in the adult epidermis and hair follicles as well (Figure S1B) (Joost et al., 2020). Bulk RNA-seq of P4 full-thickness mouse back skin and of epidermal sheets from 3-week-old, 6-month-old, and 22-month-old mice shows that among AP-2 transcription family members, AP-2α is the most highly expressed in the adult epidermis throughout life (Figure 1A). In 3-week-old back skin epidermis, ∼53% of AP-2 transcripts are for AP-2α, ∼22% for AP-2β, ∼21% for AP-2γ, and ∼4% for AP2ε (AP-2δ is not expressed in the epidermis) (Figure 1A). During the early postnatal phase (P4), AP-2β is the second most highly expressed AP-2 family member in the skin, followed by AP-2γ, whereas AP-2ε is expressed only at low levels (Figure 1A). In the epidermis of 6- or 22-month-old mice, AP-2β expression decreases compared to mice that are 3 weeks of age (Figure 1A). Notably, inactivation of AP-2α and AP-2β in the epidermis does not result in a compensatory increase in AP-2γ or AP-2ε expression (<lg2FC) (Figures 1A and 1B). Inactivation of AP-2α does not result in AP-2β upregulation in the 3-week-old epidermis and *vice versa* as well (Figure 1B).

**Figure 1:**
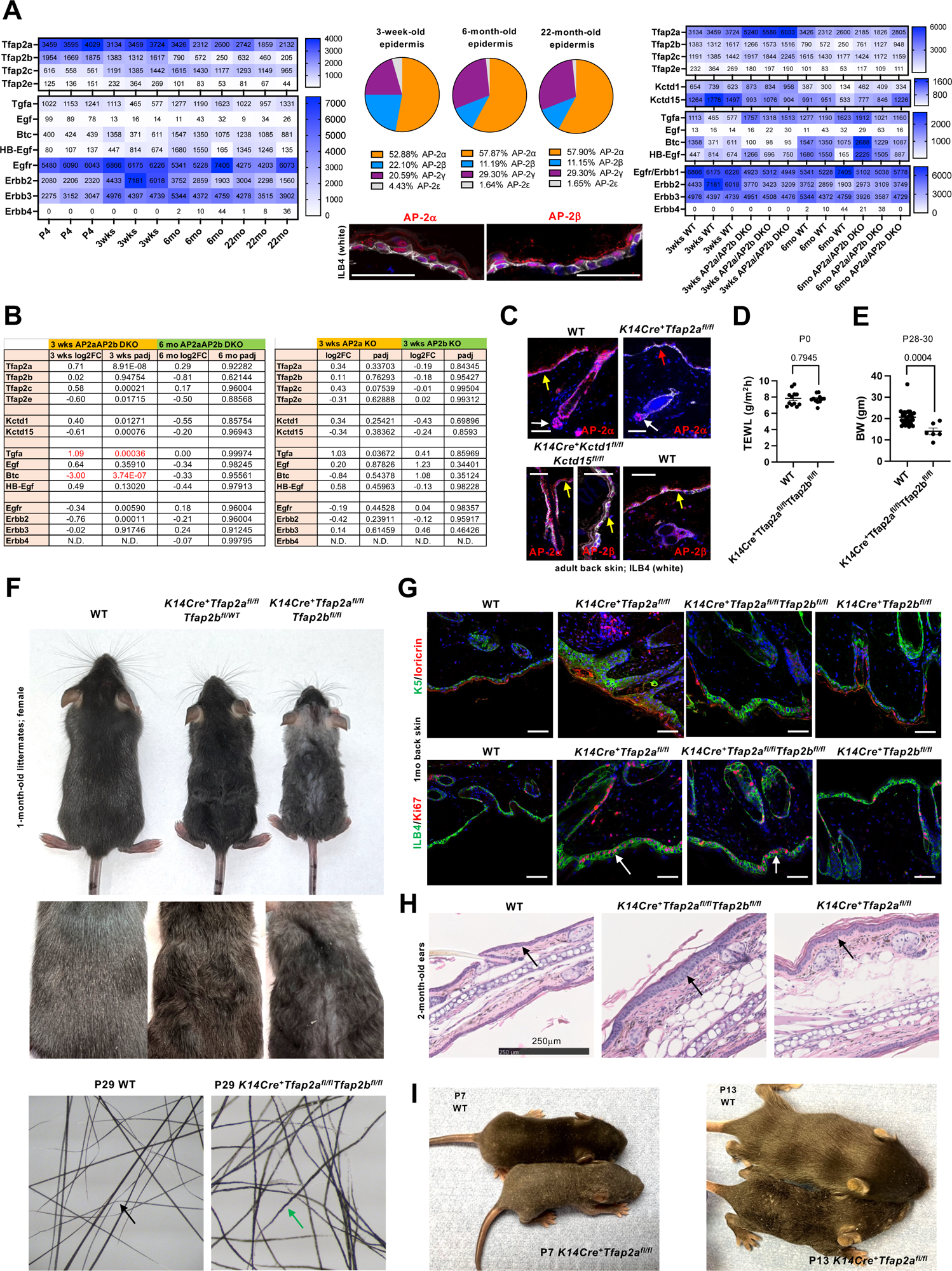
AP–2α is the most abundant AP-2 family member in the epidermis. A. Left: Heatmap shows RNA-seq transcript levels of AP-2, EGFR, and EGF family members in P4 skin and the epidermis of 3-week-old, 6-week-old, and 22-month-old WT mice. Middle: Pie charts show relative contributions of AP-2 family members to overall AP-2 transcript levels in epidermis of 3-week-old, 6-week-old, and 22-month-old WT mice. Immunolabeling of adult mouse back skin shows that AP–2α and AP–2β are located in the nuclei of basal and suprabasal keratinocytes. Isolectin B4-DyLight649 (white). Scale bars, 50 mm. Right: Effects of inactivation of AP–2α/AP–2β in keratinocytes on transcript levels of AP-2, EGFR, and EGF family members in 3-week-old and 6-month-old epidermis. B. Left: Effects of inactivation of AP–2α/AP–2β in keratinocytes on transcript levels of AP-2, EGFR, KCTD1 and KCTD15 (regulators of AP-2 activity), and EGF family members in 3-week-old and 6-month-old epidermis. Right: Values shown for 3-week-old epidermis lacking AP–2α or AP–2β in keratinocytes. Log2FC and adjusted p-values (padj) are shown. C. Immunolabeling for AP–2α and AP–2β shows overlapping expression in adult keratinocytes. Back skin of adult WT mice shown. Yellow arrows: interfollicular epidermis. the dermal papilla is not targeted by K14Cre and AP–2α is maintained in the dermal papilla of *K14Cre^+^Tfap2a^fl/fl^* mice [white arrow] that lack AP–2α immunolabeling in keratinocytes [red arrow]). Inactivation of Kctd1 and Kctd15 function in keratinocytes does not affect nuclear localization of AP–2α and AP–2β. Scale bars, 50 mm. D. Transepidermal water loss (TEWL in g/m^2^h) measurements of back skin in newborn (P0) *K14Cre^+^Tfap2a^fl/fl^Tfap2b^fl/fl^* mice and control littermates. Mean ± SEM; p-value shown for two-sided t-test. E. Body weight (BW) of male *K14Cre^+^Tfap2a^fl/fl^Tfap2b^fl/fl^* mice and control littermates at P28-P30. Mean ± SEM; p-value shown for two-sided t-test. F. Gender- and age-matched 1-month-old *K14Cre^+^Tfap2a^fl/fl^Tfap2b^fl/fl^*mice, *K14Cre^+^Tfap2a^fl/fl^Tfap2b^fl/WT^* mice, and control littermates shown. Hairs show irregular morphology with kinks, bends, and abnormal hair shaft thickness in P29 *K14Cre^+^Tfap2a^fl/fl^Tfap2b^fl/fl^* mice (green arrow). Typical zigzag hairs, seen in WT controls (black arrow), are not observed in *K14Cre^+^Tfap2a^fl/fl^Tfap2b^fl/fl^*mice. G. Top: Immunolabeling for keratin 5 (K5; green) and loricrin (red) in back skin of 1-month-old *K14Cre^+^Tfap2a^fl/fl^Tfap2b^fl/fl^* mice, *K14Cre^+^Tfap2a^fl/fl^* mice, *K14Cre^+^Tfap2b^fl/fl^* mice, and control mice. Bottom: Immunolabeling for Ki67 (red) and labeling with isolectin B4-FITC (green) in the same specimens. Arrows indicate Ki67^+^ cells. Scale bars, 50 mm. H. Representative images from 2-month-old ear skin of *K14Cre^+^Tfap2a^fl/fl^Tfap2b^fl/fl^* mice, *K14Cre^+^Tfap2a^fl/fl^* mice, and control mice. Arrows indicate epidermis. Scale bar, 250 mm. I. P7 and P13 *K14Cre^+^Tfap2a^fl/fl^* mice and control littermates are shown. AP–2α inactivation in keratinocytes results in a postnatal growth retardation, delayed hair growth and abnormal morphology of hairs and whiskers.

Thus, the lack of AP-2α and/or AP-2β in keratinocytes is not compensated for by upregulation of expression of other AP-2 family members in these cells. Efficient K14Cre-mediated functional inactivation of AP-2α and AP-2β in keratinocytes occurs through Cre-mediated excision of floxed exons 5 and 6 of *Tfap2a* in *K14Cre^+^Tfap2a^fl/fl^* mice and of exon 6 of *Tfap2b* in *K14Cre^+^Tfap2b^fl/fl^*mice, which are required for DNA binding of these transcription factors (Figure S2) (Lamontagne et al., 2022).

It has previously been reported that AP-2α represses *Egfr* expression in the epidermis, demonstrated by increased *Egfr* transcript levels in neonatal skin of *K14Cre^+^Tfap2a^fl/fl^*mice, and that loss of AP-2α may lead to hyperproliferation due to increased EGFR signaling (Wang et al., 2006). Notably, our RNA-seq data of isolated back skin epidermis did not show upregulation of transcripts of *Egfr* or other EGFR family members as a consequence of AP-2α and/or AP-2β inactivation in the epidermis when assessed at 3 weeks or 6 months of age (Figures 1A and 1B). Of the EGF family ligands, expression of *Tgfa* was upregulated and of *Btc* downregulated in 3-week-old epidermis of *K14Cre^+^Tfap2a^fl/fl^Tfap2b^fl/fl^* mice (Figures 1A and 1B). Expression of AP-2α and AP-2β in keratinocytes was not dependent on KCTD1 or KCTD15, which have been reported to inhibit AP-2α and AP-2β activity *in vitro* (Figure 1C) (Ding et al., 2009; Hu et al., 2020; Smaldone et al., 2019; Zarelli and Dawid, 2013).

Immunolabeling detected nuclear AP-2α and AP-2β expression in basal and suprabasal keratinocytes of epidermis and hair follicles (Figures 1A and 1C). In addition to keratinocytes, immunolabeling was also detected in sebocytes and dermal papillae, demonstrating an overlapping expression pattern of AP-2α and AP-2β (Figure 1C) (Lamontagne et al., 2022). For example, immunolabeling of skin of *K14Cre^+^Tfap2a^fl/fl^*mice with an anti-AP-2α antibody showed loss of AP-2α protein in keratinocytes of the epidermis and hair follicle, whereas the dermal papilla that is not targeted by the K14Cre maintains AP-2α immunolabeling (Figure 1C). The overlapping expression pattern of AP-2α and AP-2β is consistent with the observation that these transcription factors can both homodimerize and heterodimerize and have a high homology (Eckert et al., 2005; Knight et al., 2005). This suggests that functional redundancy between AP-2α and AP-2β exists in the skin. Thus, we hypothesized that inactivation of AP-2α in keratinocytes (and thereby reduction of AP-2 transcripts in the adult epidermis by ∼50%) would result in a skin phenotype that would be even more pronounced in mice lacking both AP-2α and AP-2β in keratinocytes (∼75% reduction of AP-2 transcripts), whereas loss of AP-2β function alone (reduction of <25% of AP-2 transcripts) may be compensated for by the presence of the other AP-2 family members. Analysis of *K14Cre^+^Tfap2b^fl/fl^* mice, *K14Cre^+^Tfap2a^fl/fl^* mice, and *K14Cre^+^Tfap2a^fl/fl^Tfap2b^fl/fl^*mice allows us, therefore, to assess the consequences of a progressive overall reduction of AP-2 activity in the epidermis for the manifestation of skin phenotypes.

### AP-2α and AP-2β regulate epidermal proliferation, differentiation, and hair morphogenesis

To assess the functions of AP-2α and AP-2β in the epidermis and potential compensation effects between these two AP-2 family members, we generated mice that lack AP-2α and/or AP-2β in the epidermis (*K14Cre^+^Tfap2a^fl/fl^* mice, *K14Cre^+^Tfap2b^fl/fl^*mice, *K14Cre^+^Tfap2a^fl/fl^Tfap2b^fl/fl^* mice). We also inactivated AP-2α and/or AP-2β in postnatal dermal papilla cells (*CorinCre^+^Tfap2a^fl/fl^*mice, *CorinCre^+^Tfap2b^fl/fl^* mice, *CorinCre^+^Tfap2a^fl/fl^Tfap2b^fl/fl^*mice). Corin-Cre activity occurs only in the first week postnatally (Enshell-Seijffers et al., 2010), and the inactivation of AP-2α and/or AP-2β in dermal papilla cells postnatally did not result in morphological skin or hair abnormalities (data not shown). In contrast, K14Cre-mediated excision of floxed alleles occurs early during skin morphogenesis (∼E14.5) (Dassule et al., 2000). Notably, in contrast to the reported facial abnormalities in mice with inactivation of AP-2α and AP-2β in the surface ectoderm at ∼E8.5 (using Crect transgenic mice) (Van Otterloo et al., 2022), *K14Cre^+^Tfap2a^fl/fl^Tfap2b^fl/fl^* mice with inactivation of AP-2α and AP-2β at this later stage did not have craniofacial abnormalities. Thus, craniofacial abnormalities in *Crect^+^Tfap2a^fl/fl^Tfap2b^fl/fl^*mice are a consequence of the loss of AP-2α/AP-2β functions in the surface ectoderm that occurs between E8.5 (Crect activity onset) and E14.5 (K14Cre activity onset).

Inactivation of AP-2α and AP-2β in keratinocytes did not affect transepidermal water loss (TEWL) in newborn mice (TEWL for P0 *K14Cre^+^Tfap2a^fl/fl^Tfap2b^fl/fl^*mice shown in Figure 1D). We observed a postnatal growth retardation in *K14Cre^+^Tfap2a^fl/fl^*mice that was even more apparent in age- and gender-matched *K14Cre^+^Tfap2a^fl/fl^Tfap2b^fl/fl^* mice (Figures 1E and 1F). Inactivation of AP–2α in keratinocytes resulted also in wavy and irregularly curved hairs and whiskers, phenotypes apparent particularly during weeks 2 to 6 of life (Figure 1F). *K14Cre^+^Tfap2b^fl/fl^* mice did not show morphological skin or hair abnormalities. However, combined inactivation of AP-2α and AP-2β in keratinocytes (*K14Cre^+^Tfap2a^fl/fl^Tfap2b^fl/fl^*mice) resulted in a more severe skin/hair phenotype with increased hair loss than seen in *K14Cre^+^Tfap2a^fl/fl^* mice (Figure 1F). This suggests that (1) lack of AP–2β can be compensated for by the more abundant AP-2α in keratinocytes, (2) loss of AP-2α cannot be compensated for by AP–2β, and (3) AP-2α and AP–2β have partially interchangeable functions in keratinocytes and the combined inactivation has a greater effect than inactivation of AP-2α alone.

Morphological hair shaft abnormalities were observed in *K14Cre^+^Tfap2a^fl/fl^* mice, including kinks and irregular changes in the diameter of the hair shaft and the absence of typical zigzag hairs (Figures 1F and S3A). The wavy hair phenotype resembled macroscopically that seen in *Tgfa^-/-^*mice and *Egfr* mutants (Luetteke et al., 1993; Schneider et al., 2008). However, the wavy hair coat phenotype in *K14Cre^+^Tfap2a^fl/fl^* mice was particularly apparent during hair morphogenesis and the first hair cycle and became less apparent after ∼6 weeks of age. In contrast, the wavy hair phenotype persists in *Tgfa^-/-^* mice throughout life (Figures S3B and S3C). Moreover, hair morphological defects were distinct between *K14Cre^+^Tfap2a^fl/fl^* mice (irregular hair shaft diameter and kinks in the hairs) and *Tgfa^-/-^* mice (wavy hairs) (Figures 1F and S3A-C). Skin of *Tgfa^-/-^* mice did not show thickening of the epidermis or an inflammatory infiltrate even at an advanced age (Figure S3C). Thus, the hair phenotype is distinct between *K14Cre^+^Tfap2a^fl/fl^*mice and *Tgfa^-/-^* mice. Consistent with this observation, *K14Cre^+^Tfap2a^fl/fl^*mice did not show a reduction in epidermal *Tgfa* expression (Figure 1A).

Immunolabeling for the basal keratinocyte marker keratin 5 (K5) and the cornified layer protein loricrin showed maintained epidermal stratification with an expanded K5^+^ basal cell layer in both *K14Cre^+^Tfap2a^fl/fl^* mice and *K14Cre^+^Tfap2a^fl/fl^Tfap2b^fl/fl^*mice (Figure 1G). Lack of AP–2α or both AP–2α/AP–2β, but not of AP–2β alone, resulted in an increase in proliferating Ki67^+^ basal keratinocytes, accompanied by a thickening of the epidermis (acanthosis), consistent with keratinocyte hyperproliferation (Figures 1G and 1H). This is similar to the previously reported epidermal hyperproliferation that was already observed in P3 *K14Cre^+^Tfap2a^fl/fl^* mice (Wang et al., 2006) and suggests that AP–2α is important for the switch from proliferation towards terminal differentiation of keratinocytes.

*K14Cre^+^Tfap2a^fl/fl^* mice showed a delay in hair growth in the first few weeks after birth: at P7, hair growth appeared delayed and reduced compared to WT littermates; at P13, hair appeared shorter and irregularly curved in *K14Cre^+^Tfap2a^fl/fl^* mice (Figure 1I). Whiskers of *K14Cre^+^Tfap2a^fl/fl^* mice were also irregularly curved (Figure S3A). The hair abnormalities in *K14Cre^+^Tfap2a^fl/fl^*mice are consistent with a keratinocyte differentiation defect as a consequence of AP–2α loss in these mice.

### Lack of AP–2α and AP–2β in keratinocytes leads to a skin inflammation

Starting already at ∼3 weeks of age, *K14Cre^+^Tfap2a^fl/fl^Tfap2b^fl/fl^*mice developed a skin inflammation with erythema and hyperkeratotic scale (Figure 2A). Histologically, P24 back skin epidermis of these *K14Cre^+^Tfap2a^fl/fl^Tfap2b^fl/fl^*mice showed acanthosis, hyperkeratosis, an inflammatory infiltrate in the dermis, as well as subcorneal neutrophilic microabscesses (Figure 2B). Immunolabeling for the neutrophil marker Ly6G confirmed the accumulation of neutrophils in the dermis, epidermis, and subcorneal microabscesses of these mice (Figure 2C). In addition, F4/80^+^ macrophages, CD4^+^ T cells, and Foxp3^+^ Treg cells accumulate at sites of skin inflammation in *K14Cre^+^Tfap2a^fl/fl^Tfap2b^fl/fl^*mice (Figure 2C). The interfollicular epidermis of these mice showed an increased proportion of proliferating keratinocytes (Ki67^+^) coupled with a strong increase of the alarmin keratin Krt6a, demonstrating hyperproliferation with aberrant keratinocyte differentiation (Figure 2C). Similar to control skin, nuclear Lef1 immunolabeling, a marker for activation of b-catenin signaling, was localized only in the lower hair follicle but not in the interfollicular epidermis (Figure 2C).

**Figure 2:**
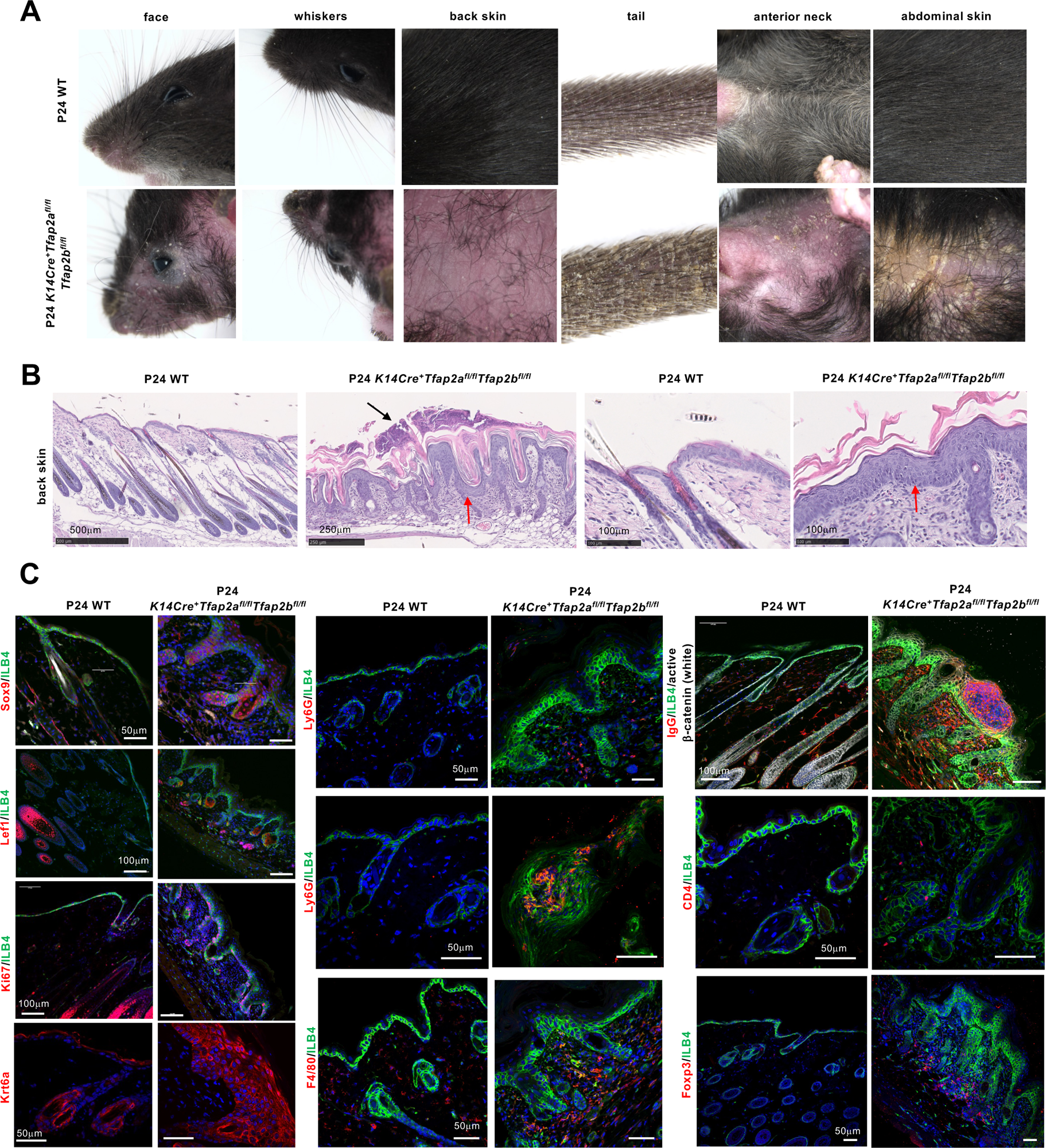
Lack of AP–2α and AP–2β in keratinocytes leads to a skin inflammation. A. Representative images of P24 WT and *K14Cre^+^Tfap2a^fl/fl^Tfap2b^fl/fl^* littermate mice are shown. Lack of AP–2α and AP–2β in keratinocytes leads to a widespread skin inflammation with erythema, hyperkeratotic scale, and hair loss. Images of the head, upper back, tail, and anterior neck are shown. Whiskers and fur hair are abnormal in *K14Cre^+^Tfap2a^fl/fl^Tfap2b^fl/fl^* mice. B. H&E images from the back skin of P24 WT and *K14Cre^+^Tfap2a^fl/fl^Tfap2b^fl/fl^* littermate mice shown in A. P24 *K14Cre^+^Tfap2a^fl/fl^Tfap2b^fl/fl^*back skin shows neutrophilic subcorneal microabscesses (black arrow) and acanthosis (red arrow). Scale bars shown in images. C. Immunolabeling of skin section from back skin of mice shown in A. and B. ILB4 labels keratinocytes. Immunolabelings with antibodies against Ly6G (neutrophils), CD4 (T helper cells), Foxp3 (Treg cells), F4/80 (macrophages), Sox9 (hair follicle stem cells), Lef1 (b-catenin signaling), active β-catenin, IgG antibodies, Ki67 (proliferation marker), and Krt6a are shown. White arrows show subcorneal microabscesses with neutrophils (Ly6G^+^ cells). Scale bars shown in images.

In contrast, *K14Cre^+^Tfap2a^fl/fl^* mice developed a similar skin inflammation only later in life, mainly after 6 months of age (Figure 3A). *K14Cre^+^Tfap2a^fl/fl^* mice developed a skin inflammation first on the face (periorbital skin and whisker area) and anterior neck skin, followed by involvement of the back skin (Figure 4A). At that age, these mice often also developed epidermal inclusion cysts on the lower abdomen (Figure 3A). Comparison of age- and gender-matched littermates demonstrates that inactivation of both AP–2α and AP–2β results in an earlier onset of a skin inflammation compared to the loss of AP–2α alone, consistent with an overall more severe phenotype in *K14Cre^+^Tfap2a^fl/fl^Tfap2b^fl/fl^* mice than in *K14Cre^+^Tfap2a^fl/fl^* mice (Figure 3B). In contrast, lack of AP–2β in keratinocytes (*K14Cre^+^Tfap2b^fl/fl^* mice) did not result in skin inflammation or other morphological skin abnormalities even in mice greater than 12 months of age.

**Figure 3:**
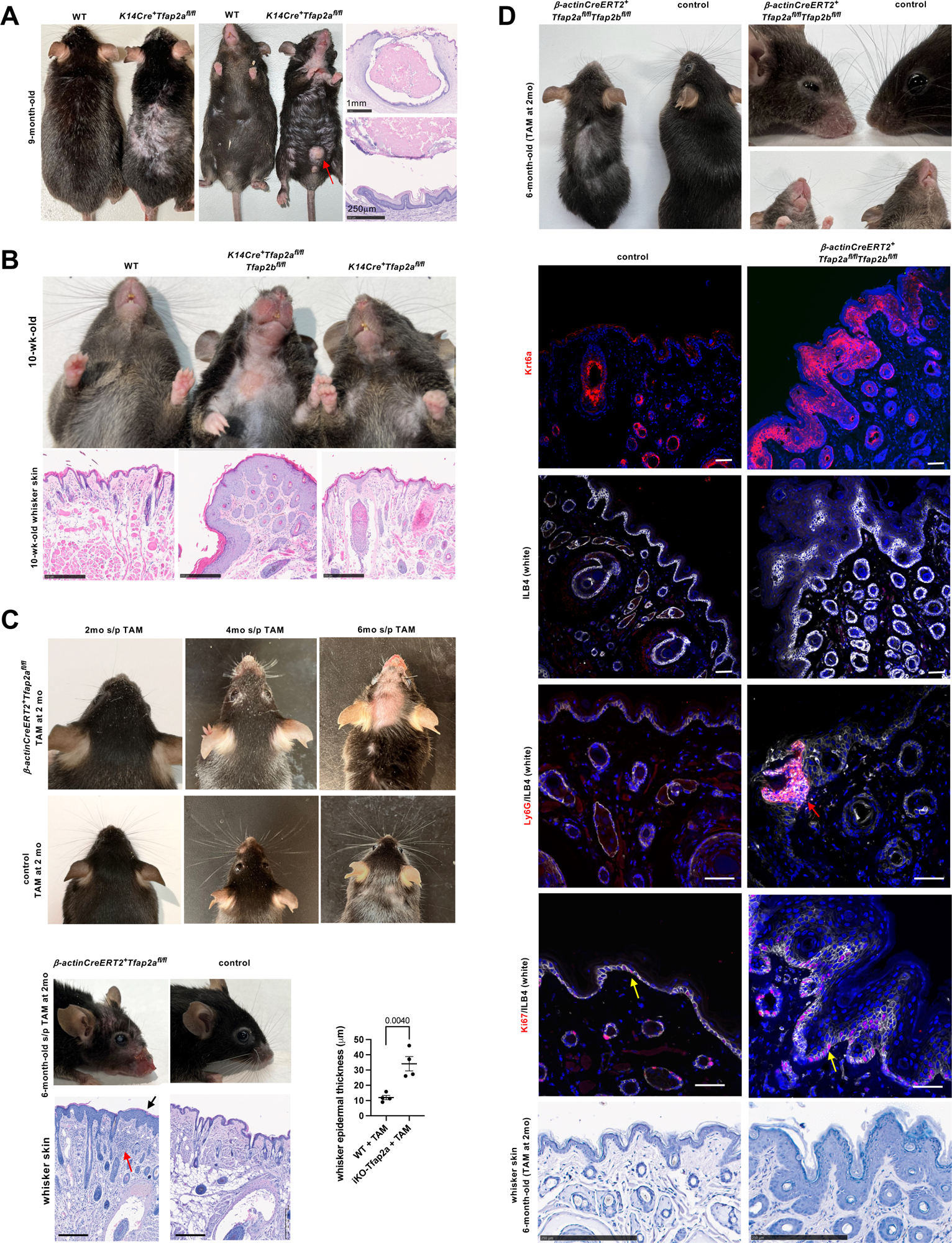
Skin inflammation worsens with age progression in *K14Cre^+^Tfap2a^fl/fl^* mice and in mice with inducible inactivation of AP–2α and AP–2α/AP–2β in the adult. A. 9-month-old *K14Cre^+^Tfap2a^fl/fl^* mice show a skin inflammation with hair loss affecting particularly the face, anterior neck, and back. Epidermoid cysts are observed in male mice on the abdomen (red arrow; H&E images). Scale bars: top 1mm, bottom 250 mm. B. 10-week-old *K14Cre^+^Tfap2a^fl/fl^Tfap2b^fl/fl^* mice show a severe skin inflammation affecting the whisker area and anterior neck, whereas at this time littermate *K14Cre^+^Tfap2a^fl/fl^* mice have not developed a severe skin inflammation yet. H&E images of whisker skin show acanthosis in *K14Cre^+^Tfap2a^fl/fl^Tfap2b^fl/fl^*mice that is more pronounced than in *K14Cre^+^Tfap2a^fl/fl^*mice at 10 weeks of age. Scale bars, 250 mm. C. Top: Inducible inactivation of AP–2α at 2 months of age (*β-actinCreERT2^+^Tfap2a^fl/fl^*mice injected with TAM at 2 months of age) and assessed 2, 4, and 6 months later. A progressive worsening of the skin inflammation with loss of whiskers is observed in mice with induced inactivation of AP–2α (images of head are shown; littermate Cre-negative controls show no skin changes). Bottom: Six months after inactivation of AP–2α at 2 months of age a severe skin inflammation is seen in the whisker area with acanthosis (black arrow) and a dermal inflammatory infiltrate (red arrow). Scale bars, 250 mm. Graph shows significant thickening of the epidermis. Mean ± SEM; p-value shown for two-sided t-test. D. Induced inactivation of AP–2α and AP–2β at 2 months of age (*β-actinCreERT2^+^Tfap2a^fl/fl^ Tfap2b^fl/fl^*mice injected with TAM at 2 months of age) and assessed 4 months later shows skin inflammation in the whisker area and inflammation and hair loss of the back skin. Immunolabelings for Krt6a, Ly6G (red arrow), and Ki67 (yellow arrows) of whisker skin of these mice. ILB4 labels keratinocytes. Scale bars, 50 mm. Corresponding H&E images of these mice are shown as well (scale bars, 250 mm).

**Figure 4:**
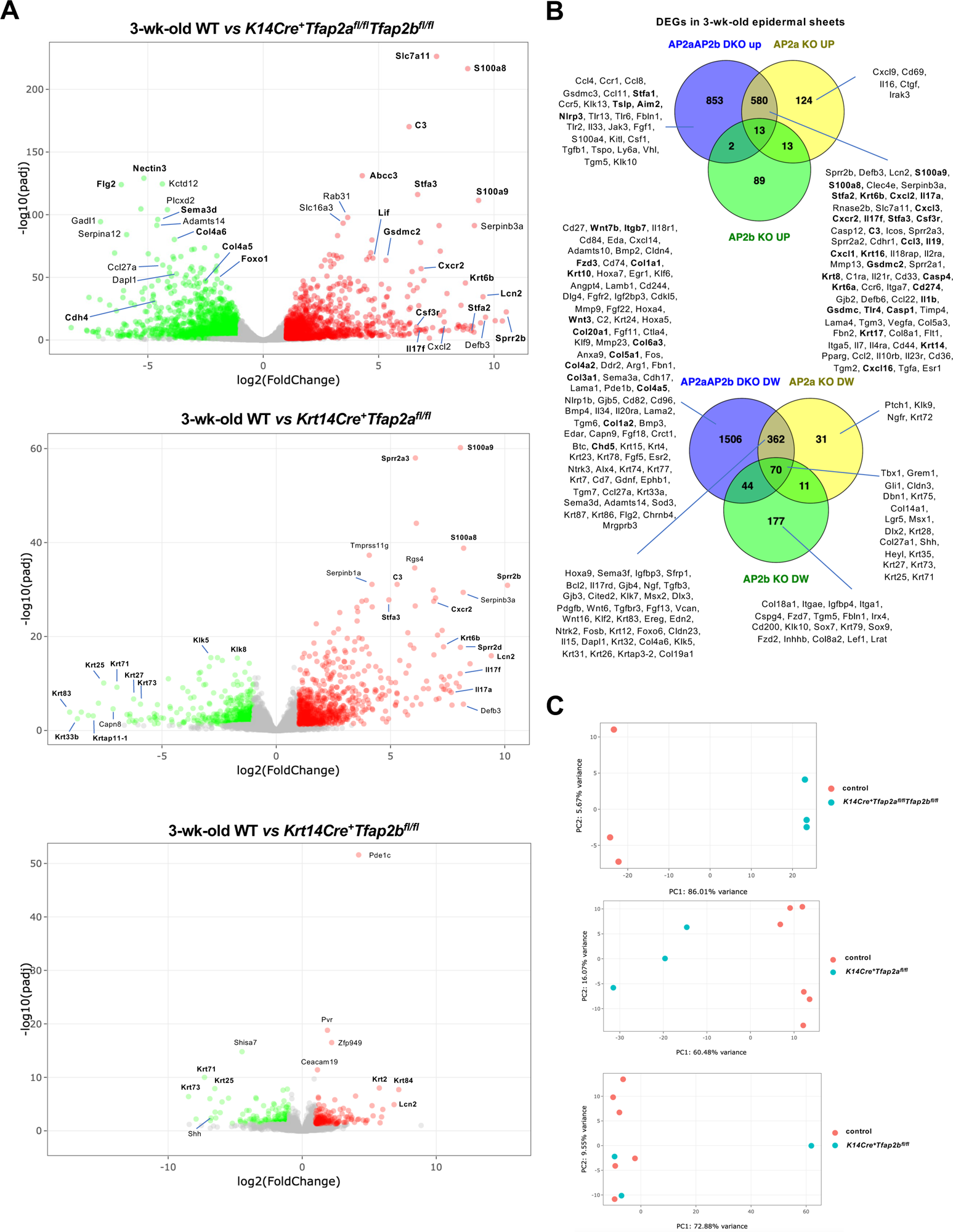
Transcriptome of epidermis lacking AP–2α and/or AP–2β in keratinocytes of 3-week-old mice. A. Volcano plots of RNA-seq data show DEGs expressed in the back skin epidermis of 3-week-old *K14Cre^+^Tfap2a^fl/fl^* mice, *K14Cre^+^Tfap2b^fl/fl^* mice, and *K14Cre^+^Tfap2a^fl/fl^Tfap2b^fl/fl^* mice compared to WT controls. Significantly upregulated DEGs in mutant mice are in red and downregulated DEGs are in green. Selected genes are shown. B. Top: Venn diagrams show the overlap of DEGs found to be upregulated in the back skin epidermis of 3-week-old *K14Cre^+^Tfap2a^fl/fl^* mice (yellow), *K14Cre^+^Tfap2b^fl/fl^* mice (green), and *K14Cre^+^Tfap2a^fl/fl^Tfap2b^fl/fl^* mice (blue) compared to WT controls. Selected upregulated DEGs are shown. Selected downregulated DEGs are shown below. C. Principal component (PC) analysis between these groups of mice for their RNA-seq data of back skin epidermis.

### Inducible inactivation of AP–2α and AP–2α/AP–2β

Inactivation of AP–2α in the adult mice (tamoxifen [TAM] administration to *b-actinCreERT2^+^Tfap2a^fl/fl^* mice at 8 weeks of age and assessment of these mice up to 9 months of age) also resulted in epidermal acanthosis and a skin inflammation that became more severe with age progression (Figure 3C). Similarly as observed in *K14Cre^+^Tfap2a^fl/fl^* mice, the skin inflammation manifested first in the whisker and periorbital area (Figure 3C). These mice lost their whiskers but did not develop curly fur hairs, consistent with our observation that the curly hair phenotype in *K14Cre^+^Tfap2a^fl/fl^* mice presents in the early postnatal period and wanes after ∼6 weeks of age (Figure 3C). Thus, the inactivation of AP–2α in the adult leads to a skin inflammation that resembles the phenotype observed in *K14Cre^+^Tfap2a^fl/fl^* mice, demonstrating a critical role of AP–2α for skin homeostasis in the adult.

Inducible inactivation of both AP–2α and AP–2β (tamoxifen [TAM] administration to *β-actinCreERT2^+^Tfap2a^fl/fl^Tfap2b^fl/fl^*mice at 8 weeks of age) resulted in a more severe skin inflammation with an earlier onset compared to those with inducible inactivation of AP–2α alone (Figure 3D), similarly as observed when comparing age-matched *K14Cre^+^Tfap2a^fl/fl^*mice and *K14Cre^+^Tfap2a^fl/fl^Tfap2b^fl/fl^* mice. Analysis of whisker skin in 6-months-old mice in which AP–2α and AP–2β were inactivated at 2 months of age showed similar changes as seen in skin of *K14Cre^+^Tfap2a^fl/fl^Tfap2b^fl/fl^*mice, including acanthosis, Krt6a upregulation in keratinocytes, increased Ki67^+^ keratinocytes, and neutrophilic microabscesses in the epidermis (Figure 3D). Similarly as with *K14Cre^+^Tfap2b^fl/fl^* mice, inducible inactivation of AP–2β (tamoxifen [TAM] administration to *β-actinCreERT2^+^Tfap2b^fl/fl^*mice at 8 weeks of age) did not result in a skin inflammation.

### Transcriptome of epidermis lacking AP–2α or AP–2α/AP–2β shows activation of multiple inflammatory pathways

Next, we performed RNA-seq of 3-week-old epidermis (transition phase from telogen to anagen hair follicles) from *K14Cre^+^Tfap2a^fl/fl^* mice, *K14Cre^+^Tfap2b^fl/fl^* mice, *K14Cre^+^Tfap2a^fl/fl^Tfap2b^fl/fl^*mice, and control mice. Consistent with the lack of a skin phenotype in *K14Cre^+^Tfap2b^fl/fl^*mice, differentially expressed genes (DEGs, defined by an adjusted p-value <0.05 and a log2 fold change >1) in these mice were much fewer than those observed in *K14Cre^+^Tfap2a^fl/fl^*mice (Figures 4A, 4B, and S4A; Table S1). Combined inactivation of AP–2α and AP–2β in keratinocytes had a significant proportion of additional DEGs not observed in the epidermis lacking only AP–2α (Figures 4A and 4B). This correlates with the more severe skin phenotype in *K14Cre^+^Tfap2a^fl/fl^Tfap2b^fl/fl^* mice than in *K14Cre^+^Tfap2a^fl/fl^* mice. It also suggests that the lack of AP–2α can be partially compensated for by AP–2β. Principal component analysis (PCA) also showed a clear separation of the epidermal transcriptome of *K14Cre^+^Tfap2a^fl/fl^*mice and *K14Cre^+^Tfap2a^fl/fl^Tfap2b^fl/fl^* mice from control mice, which was not the case for the *K14Cre^+^Tfap2b^fl/fl^*group (Figure 4C).

Gene ontology (GO) analysis showed that the epidermis of 3-week-old *K14Cre^+^Tfap2a^fl/fl^Tfap2b^fl/fl^* mice had transcriptomic changes linked to changes in cell cycle and cell division (hyperproliferation phenotype), keratinocyte differentiation, and inflammatory responses (Figure 5A). Gene set enrichment analysis (GSEA) also showed changes in keratinocyte proliferation and differentiation, as well as activation of inflammation linked to neutrophils and other inflammatory pathways (Figures 5B and S4B). GSEA also linked increased oxidative phosphorylation and glycolysis with the changes observed in the epidermis of 3-week-old *K14Cre^+^Tfap2a^fl/fl^Tfap2b^fl/fl^*mice (Figures 5B and S4B).

**Figure 5:**
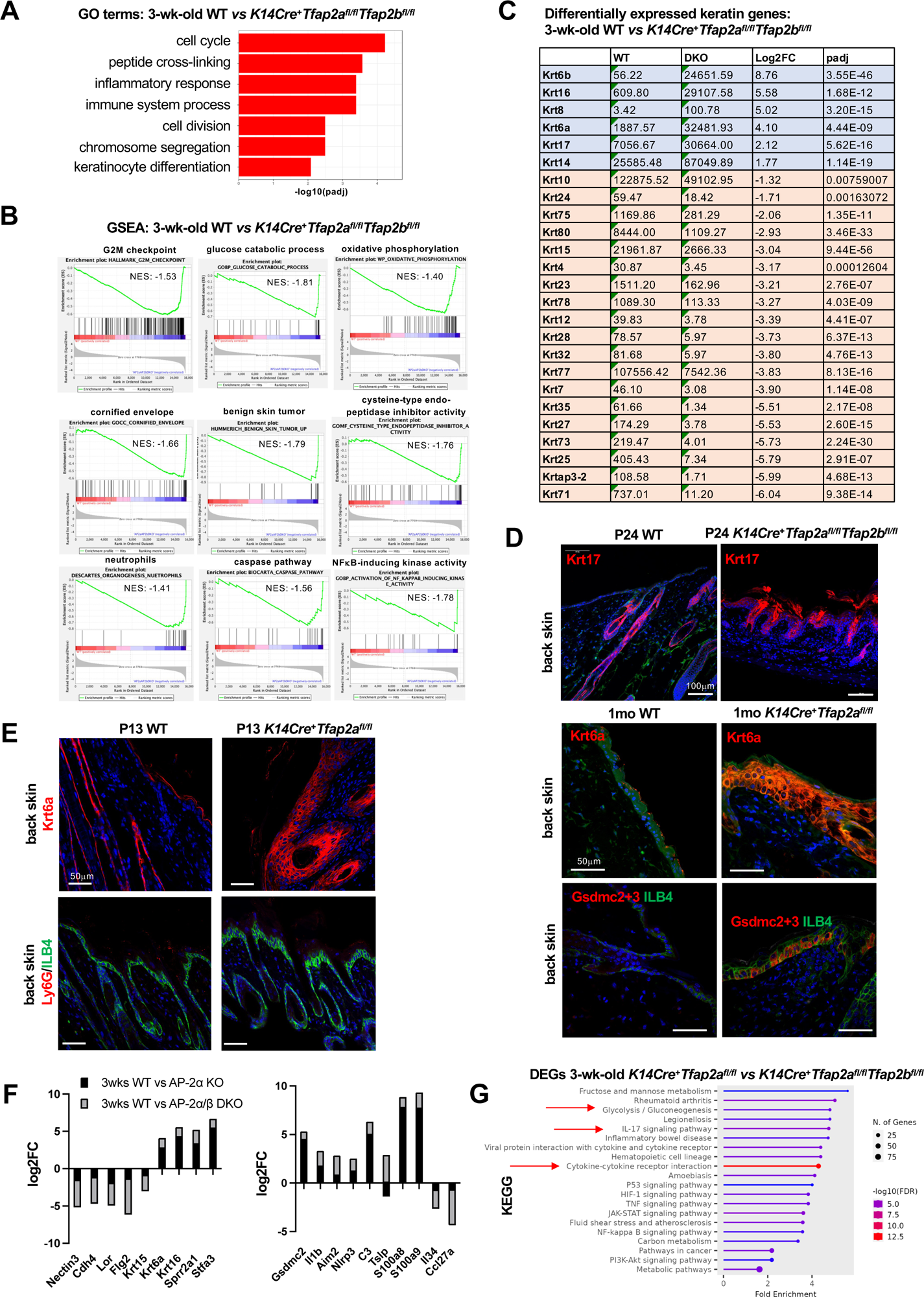
Keratinocyte hyperproliferation and terminal differentiation defect precede the neutrophilic inflammation in epidermis lacking AP–2α/AP–2β. A. GO enrichment analysis of significantly associated DEGs in back skin epidermis of 3-week-old *K14Cre^+^Tfap2a^fl/fl^Tfap2b^fl/fl^*mice. B. GSEA of RNA-seq data from back skin epidermis of 3-week-old *K14Cre^+^Tfap2a^fl/fl^Tfap2b^fl/fl^*mice compared to WT controls. Enrichment plots of selected pathways are shown. NES: normalized enrichment scores. C. Keratin genes shown that are differentially expressed in the back skin epidermis of 3-week-old *K14Cre^+^Tfap2a^fl/fl^Tfap2b^fl/fl^* mice compared to WT controls. Average transcript values, log2FC and adjusted p-values (padj) are shown. D. Top: Immunolabeling for Krt17 (red) in the back skin of P24 *K14Cre^+^Tfap2a^fl/fl^Tfap2b^fl/fl^*mice and WT mice. Bottom: Immunolabeling for Krt6a (red) and Gsdmc2+3 (red) in the back skin of 1-month-old *K14Cre^+^Tfap2a^fl/fl^* mice and WT mice. Scale bars: top 100 mm, bottom 50 mm. E. Already at P13, *K14Cre^+^Tfap2a^fl/fl^* mice have an acanthosis with accumulation of Krt6a in the interfollicular epidermis without accumulation of Ly6G^+^ neutrophils. Scale bars: 50 mm. F. Selected genes shown from RNA-seq data of back skin epidermis of 3-week-old *K14Cre^+^Tfap2a^fl/fl^Tfap2b^fl/fl^* mice and *K14Cre^+^Tfap2a^fl/fl^* mice relative to their age-matched WT controls. Graphs show log2FC of selected genes involved in keratinocyte adhesion and differentiation or skin inflammation. G. KEGG functional pathway analyses of DEGs that are up- or downregulated in the epidermis of both 3-week-old *K14Cre^+^Tfap2a^fl/fl^Tfap2b^fl/fl^*mice and *K14Cre^+^Tfap2a^fl/fl^* mice. Arrows indicate increased activation of inflammatory pathways, including IL-17 signaling, and other selected key pathways.

Epidermis of 3-week-old *K14Cre^+^Tfap2a^fl/fl^*mice and *K14Cre^+^Tfap2a^fl/fl^Tfap2b^fl/fl^* mice, compared to control mice, showed upregulation of keratins Krt6, Krt16, and Krt17, as seen with hyperproliferation or inflamed skin conditions (Figures 4A, 4B, and 5C) (Wang et al., 2006). These keratins have been described as early barrier alarmins, whose upregulation contributes to keratinocyte hyperproliferation and innate immune activation (Zhang et al., 2019). We confirmed by immunolabeling strongly increased Krt6a and Krt17 in keratinocytes of *K14Cre^+^Tfap2a^fl/fl^* mice or *K14Cre^+^Tfap2a^fl/fl^Tfap2b^fl/fl^*mice (Figures 2C, 5D, and 5E). Importantly, increased Krt6a was already observed early on in the epidermis of *K14Cre^+^Tfap2a^fl/fl^* mice (Figure 5E shows P13 mice) before an infiltration of Ly6G^+^ neutrophils, confirming that in these mice keratinocyte differentiation defects precede the inflammatory infiltrate.

In addition, expression of Krt14, expressed by basal keratinocytes, was increased as well, consistent with expansion of the basal cell compartment (Figure 5C). Krt8, normally expressed in embryonic epidermis, was highly upregulated in the epidermis of 3-week-old *K14Cre^+^Tfap2a^fl/fl^Tfap2b^fl/fl^* mice as well (Figure 5C). In contrast, keratins associated with a switch from keratinocyte proliferation to differentiation, such as the suprabasal Krt10, or other keratinocyte terminal differentiation markers (e.g., loricrin [*Lor*] or filaggrin [*Flg2*]) were downregulated in these keratinocytes (Figures 5C and 5F). These findings are consistent with a switch from terminal differentiation towards hyperproliferation when AP–2α or AP–2α/AP–2β are absent. The epidermal differentiation defect was linked to the downregulation of keratinocyte cell adhesion genes Cdh4 and Nectin3 (Figures 4A, 4B, 5F, and S4A). Notably, the more severe skin phenotype in *K14Cre^+^Tfap2a^fl/fl^Tfap2b^fl/fl^*mice compared to *K14Cre^+^Tfap2a^fl/fl^* mice correlated with a greater extent of downregulation of these keratinocyte differentiation and cell adhesion genes and the upregulation of alarmin keratin genes (Figure 5F). These findings are consistent with a more severe epidermal differentiation defect in 3-week-old *K14Cre^+^Tfap2a^fl/fl^Tfap2b^fl/fl^*mice than in age-matched *K14Cre^+^Tfap2a^fl/fl^* mice.

Several small proline-rich proteins (SPRR proteins), serpin proteins, and late cornified envelope proteins (LCE proteins), which regulate keratinocyte differentiation processes, can be strongly increased in inflammatory skin conditions (e.g., SERPINB4, SPRR2A, SERPINB3, SPRR2D, SPRR2B, SPRR2F, LCE3A) (Wang et al., 2023). Similarly, we found in epidermis of *K14Cre^+^Tfap2a^fl/fl^Tfap2b^fl/fl^*mice or *K14Cre^+^Tfap2a^fl/fl^* mice highly increased expression of several SPRR proteins (Sprr2b, Sprr2a2, Sprr2a3, Sprr2a1), serpins (Serpinb3a), or LCE proteins (Lce3d, Lce3e) (Figures 4A, 4B, and S4A). Stefin proteins (Stfa2, Stfa3) with cathepsin inhibitor activity, involved in keratinocyte differentiation, and found in the cornified cell envelope of keratinocytes, were highly upregulated in the epidermis of *K14Cre^+^Tfap2a^fl/fl^* mice and *K14Cre^+^Tfap2a^fl/fl^Tfap2b^fl/fl^* mice as well, whereas Stfa1 was upregulated only in the skin of *K14Cre^+^Tfap2a^fl/fl^Tfap2b^fl/fl^*mice (Figures 4A, 4B, and S4A).

Among the most significantly upregulated genes in the epidermis of *K14Cre^+^Tfap2a^fl/fl^* mice, and even more so in *K14Cre^+^Tfap2a^fl/fl^Tfap2b^fl/fl^* mice, were the gasdermin genes *Gsdmc* and *Gsdmc2* (Figures 4A, 4B, 5F, and S4A), whose function in keratinocytes is poorly understood. Immunolabeling with an antibody that detects Gsdmc2/Gsdmc3 showed strong immunolabeling in basal keratinocytes of the epidermis of 3-4-week-old *K14Cre^+^Tfap2a^fl/fl^* mice or *K14Cre^+^Tfap2a^fl/fl^Tfap2b^fl/fl^* mice, but not in control mice (Figures 5D and S4C). This signal likely corresponds mainly to Gsdmc2, as Gsdmc3 transcript levels were much lower than Gsdmc2 in the epidermis of these mice (Figure S4A).

Consistent with the histological presentation of a skin inflammation with neutrophilic subcorneal microabscesses, the RNA-seq data from 3-week-old epidermis of *K14Cre^+^Tfap2a^fl/fl^* mice and *K14Cre^+^Tfap2a^fl/fl^Tfap2b^fl/fl^* mice showed upregulation of cytokines linked to activation of neutrophils (Figures 5B and S4A). Th17 cytokines were upregulated in 3-week-old epidermis of *K14Cre^+^Tfap2a^fl/fl^* mice and *K14Cre^+^Tfap2a^fl/fl^Tfap2b^fl/fl^* mice as well, including Il17a, Il17f, and Il21r (Figures 4A, 4B, and S4A). KEGG analysis also showed increased IL-17 signaling pathway activation in the epidermis of these mice (Figure 5G). CCR6 was upregulated in the epidermis of *K14Cre^+^Tfap2a^fl/fl^Tfap2b^fl/fl^* mice as well (Figure S4). IL-34 expression was downregulated in the epidermis of *K14Cre^+^Tfap2a^fl/fl^Tfap2b^fl/fl^* mice (Li et al., 2014) (Figure S4A).

IL-1β, an NFκB-activating cytokine that maintains cytokine expression in Th17 cells, was upregulated as well (Figure S4A). Increased IL-1β promotes the presence of neutrophils through the production of CXCL chemokines and IL-23 pathway activation (LCN2), which were both highly increased in the epidermis of *K14Cre^+^Tfap2a^fl/fl^Tfap2b^fl/fl^*mice (Xi et al., 2022) (Figure S4A). As such, chemokine ligands Cxcl1 and Cxcl3 (chemotactic for neutrophils), Cxcl2 (produced by activated monocytes and neutrophils at sites of inflammation), the chemokine receptor Cxcr2 (receptor for Cxcl1, involved in neutrophil activation), and Csf3r (a key regulator of neutrophil proliferation and differentiation) were highly upregulated in the epidermis of *K14Cre^+^Tfap2a^fl/fl^*mice and *K14Cre^+^Tfap2a^fl/fl^Tfap2b^fl/fl^* mice (Figure S4A). These changes are reflective of a neutrophilic inflammatory process.

Treg receptor CD25 (Il2ra) expression was strongly increased in the epidermis of *K14Cre^+^Tfap2a^fl/fl^Tfap2b^fl/fl^*mice, consistent with the observed accumulation of Foxp3^+^ Tregs in the skin of these mice (Figures 2C and S4A). The dendritic cell markers CD274 (PD1-L1), CD14, and BDCA3 (THBD) were highly upregulated (Figure S4A). Elevated expression of the costimulatory molecule ICOS suggests ongoing T-cell activation (Figure S4A). A high upregulation of complement C3 in *K14Cre^+^Tfap2a^fl/fl^* mice, and an even greater increase in *K14Cre^+^Tfap2a^fl/fl^Tfap2b^fl/fl^*mice, was observed (Figures 5F and S4A).

FABP5 and S100a8, which can promote neutrophil chemotaxis and adhesion, were highly increased in the epidermis of *K14Cre^+^Tfap2a^fl/fl^Tfap2b^fl/fl^* mice (Figures 5F and S4A) (Kim et al., 2023). Increased expression of S100a8 and S100a9 was observed in the epidermis of both *K14Cre^+^Tfap2a^fl/fl^*mice and *K14Cre^+^Tfap2a^fl/fl^Tfap2b^fl/fl^* mice (Figures 5F and S4A). The metalloreductase Steap4 was also increased in the epidermis of *K14Cre^+^Tfap2a^fl/fl^Tfap2b^fl/fl^* mice (Figure S4A). We also observed a downregulation of the EGF-family ligand β-cellulin (Btc) (Figure 1A) and an upregulation of the angiogenic factor VEGF-A (Figure S4A).

DEGs that were significantly upregulated in *K14Cre^+^Tfap2a^fl/fl^Tfap2b^fl/fl^*mice but not in *K14Cre^+^Tfap2a^fl/fl^* mice included *Tslp* (enhances maturation of CD11c^+^ dendritic cells; induces the release of T-cell-attracting chemokines from monocytes), and the inflammasome pattern recognition receptors Aim2 and Nlrp3 (Figures S4A and S5B). Expression of caspase-1 and caspase-4 were upregulated already in both groups of mice (Figure S4A). Notably, Aim2 upregulation has been implicated in the epigenetic memory to inflammation in keratinocytes (Naik et al., 2017). In contrast, markers of a Th2-driven skin inflammation (e.g., IL-4, IL-5, IL-13, CCL17, or CCL18) were not increased in the epidermis of *K14Cre^+^Tfap2a^fl/fl^Tfap2b^fl/fl^*mice.

Overall, the transcriptomic changes in age-matched *K14Cre^+^Tfap2a^fl/fl^Tfap2b^fl/fl^*mice compared to *K14Cre^+^Tfap2a^fl/fl^* mice show that loss of AP–2α and AP–2β in keratinocytes results in a more severe keratinocyte differentiation defect than loss of AP–2α alone, and that this defect is associated with a more severe neutrophilic inflammatory response.

A comparison of the expression data obtained from E10.5 heads of *CrectCre^+^Tfap2a^fl/fl^Tfap2b^fl/fl^*mice (Van Otterloo et al., 2022) with the data from 3-week-old epidermis of *K14Cre^+^Tfap2a^fl/fl^Tfap2b^fl/fl^* mice showed an overlap in downregulated DEGs for the surface ectoderm markers Cxcl14, Wnt3, and Krt15 that are expressed in keratinocytes during development, but not for most other DEGs in 3-week-old epidermis of *K14Cre^+^Tfap2a^fl/fl^Tfap2b^fl/fl^*mice (Figures 6A and S6). This suggests that most changes in DEGs affecting epidermal differentiation and inflammation that lead to a skin inflammation in these mice occur after E10.5. GO analysis of DEGs obtained from E10.5 heads of *CrectCre^+^Tfap2a^fl/fl^Tfap2b^fl/fl^*mice supports an important role of AP–2α/AP–2β for proper epidermal development and a role for regulation of Wnt signaling in the skin at that developmental timepoint (Figure S5) (Van Otterloo et al., 2022). Similarly, GO analysis of multiomics data from human embryonic stem cell-derived (hESC) surface ectoderm cells (Collier et al., 2023) shows that AP–2α-dependent genes are involved primarily in the regulation of keratinocyte functions and their interaction with the extracellular matrix rather than the regulation of the expression of inflammatory mediators (Figures 6B, S7 and S8). This is consistent with our findings that inactivation of AP–2α in keratinocytes results in a primary keratinocyte differentiation defect and that the skin inflammation occurs subsequently.

**Figure 6:**
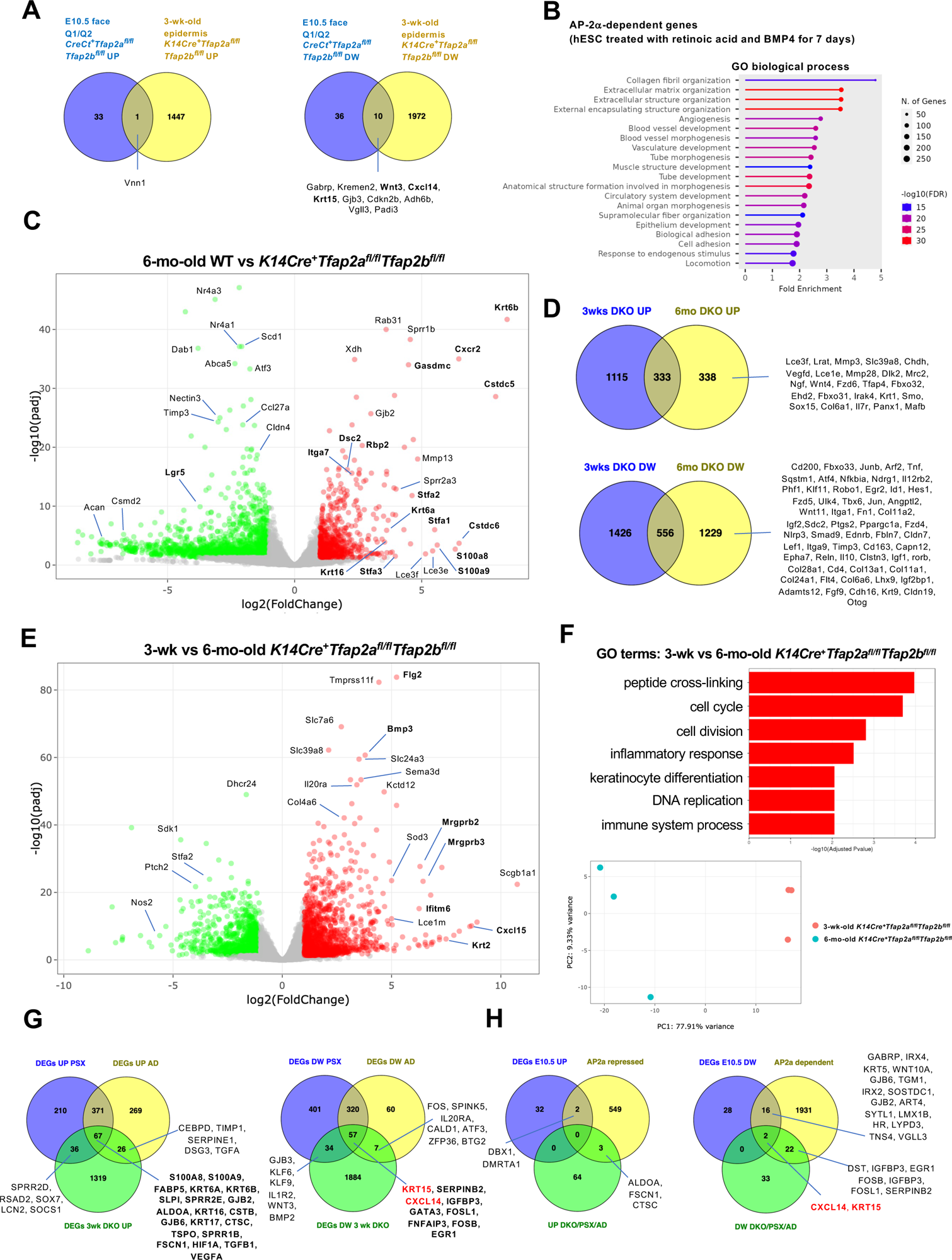
Age-dependent progressive skin inflammation due to lack of AP–2α/AP–2β. A. Venn diagrams compare DEGs in E10.5 embryonic mouse face in which AP–2α and AP–2β were inactivated at ∼E8.5 in surface ectoderm (Crect^+^*Tfap2a^fl/fl^Tfap2b^fl/fl^* mice; with a focus on epidermis-derived genes in Crect^+^*Tfap2a^fl/fl^Tfap2b^fl/fl^* mice (as described in (Van Otterloo et al., 2022)) with those of back skin epidermis from 3-week-old *K14Cre^+^Tfap2a^fl/fl^Tfap2b^fl/fl^*mice (inactivation of AP–2α and AP–2β were inactivated at ∼E14.5). B. GO biological processes associated with AP–2α-dependent genes in surface ectoderm cells derived from hESC (treated with retinoic acid and BMP4 for 7 days) (Collier et al., 2023). C. Volcano plots of RNA-seq data show DEGs expressed in the back skin epidermis of 6-month-old *K14Cre^+^Tfap2a^fl/fl^Tfap2b^fl/fl^* mice compared to WT age-matched controls. Significantly upregulated DEGs in double mutant mice in red and downregulated DEGs in green. Selected genes are shown. PCA graph for the experimental groups is shown. D. Venn diagrams show the overlap of DEGs found to be upregulated (top) or downregulated (bottom) in the back skin epidermis of 3-week-old *K14Cre^+^Tfap2a^fl/fl^Tfap2b^fl/fl^*mice (blue) compared to those of 6-months old *K14Cre^+^Tfap2a^fl/fl^Tfap2b^fl/fl^* mice (yellow). Selected genes are shown. E. Volcano plots of RNA-seq data show DEGs expressed in back skin epidermis of 3-week-old versus 6-month-old *K14Cre^+^Tfap2a^fl/fl^Tfap2b^fl/fl^*mice. Significantly upregulated DEGs in 6-month-old double mutant mice in red and downregulated DEGs in green. Selected genes are shown. F. Top: GO term analysis of DEGs expressed in back skin epidermis of 3-week-old versus 6-month-old *K14Cre^+^Tfap2a^fl/fl^Tfap2b^fl/fl^* mice. Bottom: PCA of RNA-seq data of these groups of mice. G. Venn diagrams show DEGs upregulated (left) or downregulated (right) between bulk RNA-seq data from epidermal sheets of 3-week-old *K14Cre^+^Tfap2a^fl/fl^Tfap2b^fl/fl^* mice with DEGs from scRNA-seq data of keratinocytes in human psoriasis (PSX) or atopic dermatitis (AD) (Reynolds et al., 2021). H. Venn diagrams show DEGs upregulated (left) or downregulated (right) between groups of (1) E10.5 embryonic mouse heads of Crect^+^*Tfap2a^fl/fl^Tfap2b^fl/fl^* mice [Q1/Q2], (2) AP–2α-repressed (UP) or AP–2α-dependent (DW) genes [surface ectoderm cells derived from hESC], and (3) those DEGs that are common between DKOs (epidermal sheets of 3-week-old *K14Cre^+^Tfap2a^fl/fl^Tfap2b^fl/fl^*mice) and DEGs from scRNA-seq data of keratinocytes in human psoriasis (PSX) or atopic dermatitis (AD) (Reynolds et al., 2021). Only CXCL14 and KRT15 are downregulated in all groups.

### Age-dependent progressive skin inflammation due to lack of AP–2α/AP–2β

We found that with progressive age, the skin phenotype, including skin inflammation and hair loss, worsened in *K14Cre^+^Tfap2a^fl/fl^*mice (Figure 3A) and to a more severe degree and with an earlier onset in *K14Cre^+^Tfap2a^fl/fl^Tfap2b^fl/fl^*mice, whereas no skin inflammation was observed in even >1-year-old *K14Cre^+^Tfap2b^fl/fl^*mice. Skin inflammation was also not observed even in ∼2-year-old *Tgfa^-/-^*mice. To further characterize these age-dependent changes in the epidermis, we performed RNA-seq of the epidermis of 6-month-old back skin from *K14Cre^+^Tfap2a^fl/fl^Tfap2b^fl/fl^*mice *versus* controls. Similar to changes observed in 3-week-old epidermis of these mice, we found a strong upregulation of genes involved in keratinocyte differentiation, such as cysteine-type endopeptidase inhibitors (Stfa1/2/3 and Cstdc5 and Cstdc6) or SPRR protein Sprr2a3 and Sprr2a1, and Serpinb3a. S100a8, S100a9, Gsdmc/Gsdmc2, complement C3, and keratins 6 and 16 were highly increased as well (Figures 6C, 6D, and S4A). In addition, the cell adhesion gene nectin-3 was downregulated, similarly as seen in 3-week-old epidermis of *K14Cre^+^Tfap2a^fl/fl^Tfap2b^fl/f^*mice (Figure 6C). A subset of additional DEGs involved in inflammation, keratinocyte differentiation, and interactions with the extracellular matrix, were observed in 6-month-old epidermis from *K14Cre^+^Tfap2a^fl/fl^Tfap2b^fl/fl^* mice that were not found in 3-week-old epidermis of these mice (Figures 6D-F).

### Comparison to differentially expressed genes in keratinocytes from psoriasis or atopic dermatitis lesional skin

When comparing the DEGs from the RNA-seq data derived from 3-week-old epidermis of *K14Cre^+^Tfap2a^fl/fl^Tfap2b^fl/fl^* mice with DEGs from scRNA-seq data of keratinocytes from human psoriasis and atopic dermatitis (lesional skin), a partial overlap of gene expression changes is observed (Figure 6G). Genes that were upregulated in all three groups included S100A8, S100A9, FABP5, KRT6A, KRT6B, SLPI, SPRR2E, GJB2, ALDOA, KRT16, CSTB, GJB6, KRT17, CTSC, TSPO, SPRR1B, FSCN1, HIF1A, TGFB1, and VEGFA (Figure 7G; Table S2), reflecting a proinflammatory gene expression signature in these keratinocytes. An overlap of the DEGs in the epidermis of these *K14Cre^+^Tfap2a^fl/fl^Tfap2b^fl/fl^*mice with DEGs from scRNA-seq data of keratinocytes from only lesional psoriasis skin included upregulation of SPRR2D, RSAD2, SOX7, LCN2, and SOCS1 (Figure 6G), while an overlap with DEGs from scRNA-seq data of keratinocytes from only atopic dermatitis included CEBPD, TIMP1, SERPINE1, DSG3, and TGFA (Figure 6G). Genes downregulated in all groups included KRT15, SERPINB2, CXCL14, IGFBP3, GATA3, FOSL1, FNFAIP3, FOSB, and EGR1 (Figure 6G). Notably, AP-2 transcription factor transcripts were not downregulated in keratinocytes in psoriasis or atopic dermatitis (Figure S8B).

To identify candidate genes whose expression in keratinocytes is dependent on AP–2α and diminished as an early pathogenic event that is linked to the subsequent manifestation of skin inflammation, we determined which genes are (1) downregulated in early surface ectoderm lacking AP–2α/AP2b (surface ectoderm of the face of E10.5 *CrectCre^+^Tfap2a^fl/fl^Tfap2b^fl/fl^*mice), are (2) dependent on AP–2α in differentiating human keratinocytes, and are (3) downregulated in inflammed epidermis of 3-week-old *K14Cre^+^Tfap2a^fl/fl^Tfap2b^fl/fl^*mice and in (4) keratinocytes from psoriasis and atopic dermatitis lesional skin. These analyses implicate downregulation of AP–2α-dependent gene expression of CXCL14 and KRT15 in keratinocytes as early pathogenic events that precede skin inflammation (Figure 6H).

## DISCUSSION

Conditional inactivation of AP–2α, AP–2β, or both in keratinocytes at ∼E14.5 reveals that AP–2α is a key regulator of epidermal and hair follicle differentiation, whose loss cannot be compensated for by AP–2β, whereas AP–2β loss alone does not result in hair and skin defects. Importantly, keratinocyte hyperproliferation and defects in their terminal differentiation due to loss of AP–2α precede a progressive skin inflammation. We find that AP–2α/AP–2β are not only important for hair and skin morphogenesis in the postnatal phase but that they also regulate epidermal differentiation and proliferation in the adult skin as well, as their inducible inactivation in the adult results in epidermal differentiation defects and skin inflammation with age progression.

Combined inactivation of AP–2α and AP–2β results in more severe keratinocyte differentiation defects and age-dependent skin inflammation with earlier onset than the loss of AP–2α alone, demonstrating that AP–2β can partly compensate for the loss of AP–2α and plays an overlapping role in these epidermal differentiation processes. That AP–2α has a dominant role over AP–2β for keratinocyte differentiation is explained by the higher expression of AP–2α than AP–2β in the epidermis. As both AP–2α and AP–2β have very similar DNA sequence preferences and bind as dimers to the same consensus motif, this indicates that the more severe skin phenotype in *K14Cre^+^Tfap2a^fl/fl^Tfap2b^fl/fl^*mice compared to *K14Cre^+^Tfap2a^fl/fl^* mice can be explained by the overall greater reduction in AP-2 transcription factor activity in keratinocytes, whereas *K14Cre^+^Tfap2b^fl/fl^* mice do have sufficient AP-2 activity due to the presence of the higher expressed AP–2α to prevent skin abnormalities. Notably, we have observed that loss of AP–2α does not significantly change AP–2β expression in keratinocytes and *vice versa*. Thus, our data show that the overall AP-2 transcription factor activity is diminished in the following order:

*K14Cre^+^Tfap2b^fl/fl^* mice (no skin phenotype) > *K14Cre^+^Tfap2a^fl/fl^* mice (skin phenotype) > *K14Cre^+^Tfap2a^fl/fl^Tfap2b^fl/fl^*mice (more severe skin phenotype). These findings are consistent with previous studies that provided evidence that AP-2 transcription factors can act in concert and exert redundant functions or compensate for each other’s loss (Schmidt et al., 2011; Van Otterloo et al., 2018; Van Otterloo et al., 2022; Wang et al., 2008).

That AP–2α and AP–2β can have overlapping functions and compensation mechanisms in keratinocytes is not a universal mechanism in epithelial tissues. For example, we have shown that in distal nephron epithelial cells of the kidney AP–2β and AP–2α have non-overlapping expression and function (Lamontagne et al., 2022). AP–2β is expressed in the distal nephron except the medullary collecting duct and is critical for the development and maintenance of the distal convoluted tubules, whereas AP–2α is expressed in the distal nephron mainly in the medullary collecting duct and plays a role in maintaining its structural integrity (Lamontagne et al., 2022).

Loss of AP–2α resulted in a keratinocyte hyperproliferation with increased Ki67^+^ cells and acanthosis that was associated with changes in gene expression of regulators of epidermal and hair follicle differentiation. This suggests that AP–2α functions as a major regulator for the transition of epidermal proliferation to differentiation and, when not present, results in abnormal epidermal differentiation and hyperproliferation, associated with upregulation of alarmin keratins and downregulation of specific cell adhesion genes. These changes precede a secondary skin inflammation. Notably, the comparison of the skin differentiation defects and skin inflammation in age-matched *K14Cre^+^Tfap2a^fl/fl^Tfap2b^fl/fl^*mice with those in *K14Cre^+^Tfap2a^fl/fl^* mice suggests that the extent and onset of the inflammation correlate to the extent of the keratinocyte differentiation defect.

Analysis of gene expression data in which AP–2α and AP–2β were inactivated in ectoderm at E8.5 and analyzed at E10.5, showed downregulation of genes that included Cxcl14 and Krt15, whereas many of the inflammatory genes that were upregulated in the epidermis of *K14Cre^+^Tfap2a^fl/fl^Tfap2b^fl/fl^*mice were not altered at that time point. Notably, CXCL14 and KRT15 expression in human keratinocytes is dependent on AP–2α and their expression was also downregulated in keratinocytes from the human inflammatory skin conditions psoriasis and atopic dermatitis. This suggests that CXCL14 and KRT15 downregulation may be common early pathogenic events that precede the manifestation of skin inflammation. Histologically, the acanthosis, hyperkeratosis, and subcorneal neutrophilic microabscesses observed in the skin of 3-week-old *K14Cre^+^Tfap2a^fl/fl^Tfap2b^fl/fl^*mice resemble the histologic changes seen in psoriasis. However, a comparison of the epidermal transcriptome with differentially expressed genes in keratinocytes from psoriasis and atopic dermatitis shows only a partial overlap with both inflammatory skin diseases. Furthermore, neither AP–2α nor AP–2β are downregulated in psoriasis or atopic dermatitis keratinocytes. This suggests that keratinocyte differentiation defects due to loss of AP–2α or AP–2α/AP–2β result in a secondary inflammatory keratinocyte response with partial overlap to changes seen in human inflammatory skin conditions, rather than these mice representing a mouse model of either psoriasis or atopic dermatitis.

In summary, inactivation of AP–2α or AP–2α/AP–2β transcription factors selectively in keratinocytes impairs the transition from keratinocyte proliferation to terminal differentiation and leads to a secondary skin inflammation. This suggests that a primary keratinocyte differentiation defect can be sufficient to induce a subsequent skin inflammation and that therapeutic approaches to normalize keratinocyte differentiation may inhibit or prevent certain inflammatory skin conditions. Our findings establish AP–2α/AP–2β as critical regulators of epidermal homeostasis in the adult skin.

## MATERIALS & METHODS

### Animals

*Tfap2a^fl/fl^* mice have previously been reported: Cre activity results in the removal of exons 5 and 6 that are required for DNA binding activity (Brewer et al., 2004). *Tfap2b^fl/fl^*mice have previously been described as well: Cre-mediated recombination results in removal of exon 6 which is critical for DNA binding, resulting in AP–2β without transcription factor activity (Van Otterloo et al., 2018). These mice were crossed with K14Cre^+^ mice [IMSR Cat# JAX:018964, RRID:IMSR_JAX:018694] (generating *K14Cre^+^Tfap2a^fl/fl^*mice, *K14Cre^+^Tfap2b^fl/fl^* mice, *K14Cre^+^Tfap2a^fl/fl^Tfap2b^fl/fl^*mice) or Corin-Cre mice (generating *CorinCre^+^Tfap2a^fl/fl^*mice, *CorinCre^+^Tfap2b^fl/fl^* mice, *CorinCre^+^Tfap2a^fl/fl^Tfap2b^fl/fl^*mice) (Enshell-Seijffers et al., 2010). Inducible inactivation of *Tfap2a* and *Tfap2b* in the adult was achieved in crosses with CAGGCreERT2^+^ mice (IMSR Cat# JAX:004453, RRID:IMSR_JAX:004453; referred to as *β*-actinCreERT2^+^ mice) (Hayashi and McMahon, 2002): tamoxifen [TAM] administration to *β-actinCreERT2^+^Tfap2a^fl/fl^* mice or *β-actinCreERT2^+^Tfap2a^fl/fl^Tfap2b^fl/fl^* mice; we inactivated TAM at 8-weeks and assessed these mice up to 9 months of age). We performed intraperitoneal injections of tamoxifen (TAM; T5648, Sigma Aldrich) at 6mg/40gm BW daily for 5 consecutive days, a tamoxifen administration regimen that has been shown to efficiently and uniformly activate Cre recombinase in adult tissues (Hayashi and McMahon, 2002). RNA-seq of epidermal sheets confirmed efficient Cre-mediated functional inactivation of AP–2α and AP–2β in the epidermis, which occurs through K14Cre-mediated excision of floxed exons 5 and 6 of *Tfap2a* in *K14Cre^+^Tfap2a^fl/fl^* mice and of exon 6 of *Tfap2b* in *K14Cre^+^Tfap2b^fl/fl^* mice (Supplemental Figure 3) (Lamontagne et al., 2022). For all animal studies, institutional approval was granted and international guidelines for the care and use of laboratory research animals were followed. ARRIVE guidelines for reporting animal studies were followed. TEWL was measured with a vapometer in g/m^2^h (Delfin Technologies). BW was determined in male mice.

### Immunolabeling and morphological analyses

For morphological analysis of mouse tissue, tissues were fixed in 4% paraformaldehyde. Tissues were processed and embedded in paraffin for histological analysis or immunolabeling. H&E stainings were performed according to standard protocols. Immunolabeling experiments were performed on slides that were citrate buffer-treated for heat-mediated antigen retrieval. Sections were permeabilized in 0.5% Triton X-100 and subsequently blocked with serum in which the secondary antibodies were raised. DAPI was used to stain nuclei (Thermo Fisher Scientific Cat# D3571, RRID:AB_2307445). Primary antibodies used in this study are shown in Table:

|  |  |  |
| --- | --- | --- |
| rabbit anti-AP-2β (1:200) | Cell Signaling Technology | Cat# 2509,<br>RRID:AB_2058198 |
| guinea pig anti-Krt5 (1:200) | American Research Products | 03-GP-CK5,<br>RRID:AB_1541051 |
| rabbit anti-Krt6a (1:500) | Biolegend | 905701,<br>RRID:AB_2565052 |
| rabbit anti-Col17 (1:100) | Abcam | ab184996 |
| rabbit anti-Lef1 (1:100) | Cell Signaling Technology | 2230S,<br>RRID:AB_823558 |
| rabbit anti-Ki67 (1:100) | Abcam | ab16667,<br>RRID:AB_302459 |
| rabbit anti-loricrin (1:200) | Abcam | ab85679,<br>RRID:AB_2134912 |
| rabbit anti-AP-2α (1:200) | Abcam | ab108311,<br>RRID:AB_10861200 |
| rabbit anti-Foxp3 (1:200) | Cell Signaling Technology | Cat# 12653<br>RRID:AB_2797979 |
| rabbit anti-CD4 (1:1000) | Abcam | Cat# ab183685<br>RRID:AB_2686917 |
| rabbit anti-Ly6G (1:100) | Cell Signaling Technology | Cat# 87048<br>RRID:AB_2909808 |
| rabbit-anti F4/80 | Cell Signaling Technology | Cat# 70076<br>RRID:AB_2799771 |
| rabbit anti-Krt1 (1:200) | ProteoGenix | Cat# PAA492Mu01 |
| rabbit anti-Gsdmc2/3 (1:500) | Abcam | Cat# ab229896<br>RRID:AB_2938769 |

Secondary Alexa-488/555/647 antibodies were used at a dilution of 1:200 (Thermo Fisher). Controls included stainings with no primary antibody or with IgG control primary antibodies. FITC-conjugated or DyLight649-conjugated Griffonia Simplicifolia Lectin I (GSL I) isolectin B4 (Vector Laboratories Cat# FL-1201, RRID:AB_2314663; Cat# DL-1208.5) was used at a dilution of 1:100.

### RNA-seq

For RNA-seq, tissues from 3 to 6 mice per experimental groups were used. Epidermal sheets were isolated by treatment with dispase-II for ∼1 hour at 37°C (20mg/ml) (D4693, Sigma). Skin and epidermal sheets were homogenized with a Qiagen Tissue Lyser II. RNA was isolated with Trizol reagent (Life technologies), and further purified with the Directzol RNA MiniPrep Plus kit from Zymo (R2073) and treated with DNase. Total RNA samples were quantified using Qubit 2.0 Fluorometer (Life Technologies, Carlsbad, CA, USA) and RNA integrity was checked with 4200 TapeStation (Agilent Technologies, Palo Alto, CA, USA). For RNA-seq library construction, rRNA depletion sequencing library was prepared by using QIAGEN FastSelect rRNA HMR Kit (Qiagen, Hilden, Germany). RNA sequencing library preparation uses NEBNext Ultra II RNA Library Preparation Kit for Illumina by following the manufacturer’s recommendations (NEB, Ipswich, MA, USA). Sequencing libraries were validated using the Agilent Tapestation 4200 (Agilent Technologies, Palo Alto, CA, USA), quantified using Qubit 2.0 Fluorometer (ThermoFisher Scientific, Waltham, MA, USA) as well as by quantitative PCR (KAPA Biosystems, Wilmington, MA, USA), and loaded to Illumina NovaSeq 6000 system for 150bp paired-end sequencing.

Raw sequence reads were trimmed to remove possible adapter sequences and low-quality reads using Trimmomatic v.0.36 (Bolger et al., 2014). The trimmed reads were mapped to the mouse GRCm38 reference genome using the STAR aligner v.2.5.2b (Dobin et al., 2013). Reads mapping to annotated genes were quantified using featureCounts from the Subread package v.1.5.2 (Liao et al., 2013). Differential expression analysis was performed with DESeq2(Love et al., 2014). Genes with adjusted P values < 0.05 and absolute fold changes >1 were called as differentially expressed genes (DEGs). GSEA was performed using the GSEA v4.3.2 software (https://www.gsea-msigdb.org/gsea/index.jsp). GO enrichment analyses were performed using ShinyGO 0.77 (http://bioinformatics.sdstate.edu/go/).

### Analysis of Single-Cell RNA Sequencing Data

Publicly available scRNA data of lesional and nonlesional skin from patients with atopic dermatitis (AD) and psoriasis were downloaded from this website https://developmental.cellatlas.io/diseased-skin (Reynolds et al., 2021). Cell type annotation was available from the processed h5ad file. Expression was normalized by log transformation and scaled to 10,000 transcripts using NormalizeData function in Seurat v4.4.0. Subpopulations of Keratinocytes including Differentiated_KC*, Differentiated_KC, Undifferentiated_KC*, and Proliferating_KC were merged into one Keratinocyte cluster for downstream analysis. The FindMarkers function was used to detect differentially expressed genes between two groups of cells with the following parameters: Wilcoxon rank sum test, min.pct = 0.1, logFC.threshold = 0.25. An adjusted p-value (Bonferroni Correction) cut-off < 1×10–4 was used to identify differentially expressed genes.

### Statistics

An unpaired two-tailed Student’s t-test was used for statistical analyses. P-values <0.05 were considered to be statistically significant. P-values are indicated. All graphs show mean ± SEM.

## DATA AVAILABILITY STATEMENT

### Lead Contact

Further information and requests for resources and reagents should be directed to the Lead Contact, Alexander G. Marneros.

### Materials Availability

Mouse lines generated in this study are available upon request.

### Data and Code Availability

Processed RNA-seq data from epidermis derived from mouse mutants generated in this study is provided in Table S1 and have been deposited to the GEO database (GSE248372). ScRNA-seq data of adult human skin and skin from psoriasis and atopic dermatitis have been reported (Reynolds et al., 2021). RNA-seq data from E10.5 embryonic mouse face in which AP–2α and AP–2β were inactivated at ∼E8.5 in surface ectoderm (Crect^+^*Tfap2a^fl/fl^Tfap2b^fl/fl^* mice) have been described as well (Van Otterloo et al., 2022). AP–2α-dependent or -repressed genes in surface ectoderm cells derived from hESC (treated with retinoic acid and BMP4 for 7 days) were reported (Collier et al., 2023).

## ACKNOWLEDGMENTS

This study was supported by funding from the NIH (R01DK118134, R01DK121178, R01EY033360, R01EY033709) to A.G.M.

## AUTHOR CONTRIBUTIONS

Experiments and data analysis were performed by all authors. The project was designed by A.G.M. Writing of the manuscript and figure preparations were done by A.G.M.

## Abbreviations

GSEA: gene set enrichment analysis
DEG: differentially expressed genes
TAM: tamoxifen
GO: gene ontology
E: embryonic day
P: postnatal day
FC: fold change
hESC: human embryonic stem cell

## SUPPLEMENTAL FIGURES

**Figure S1:**
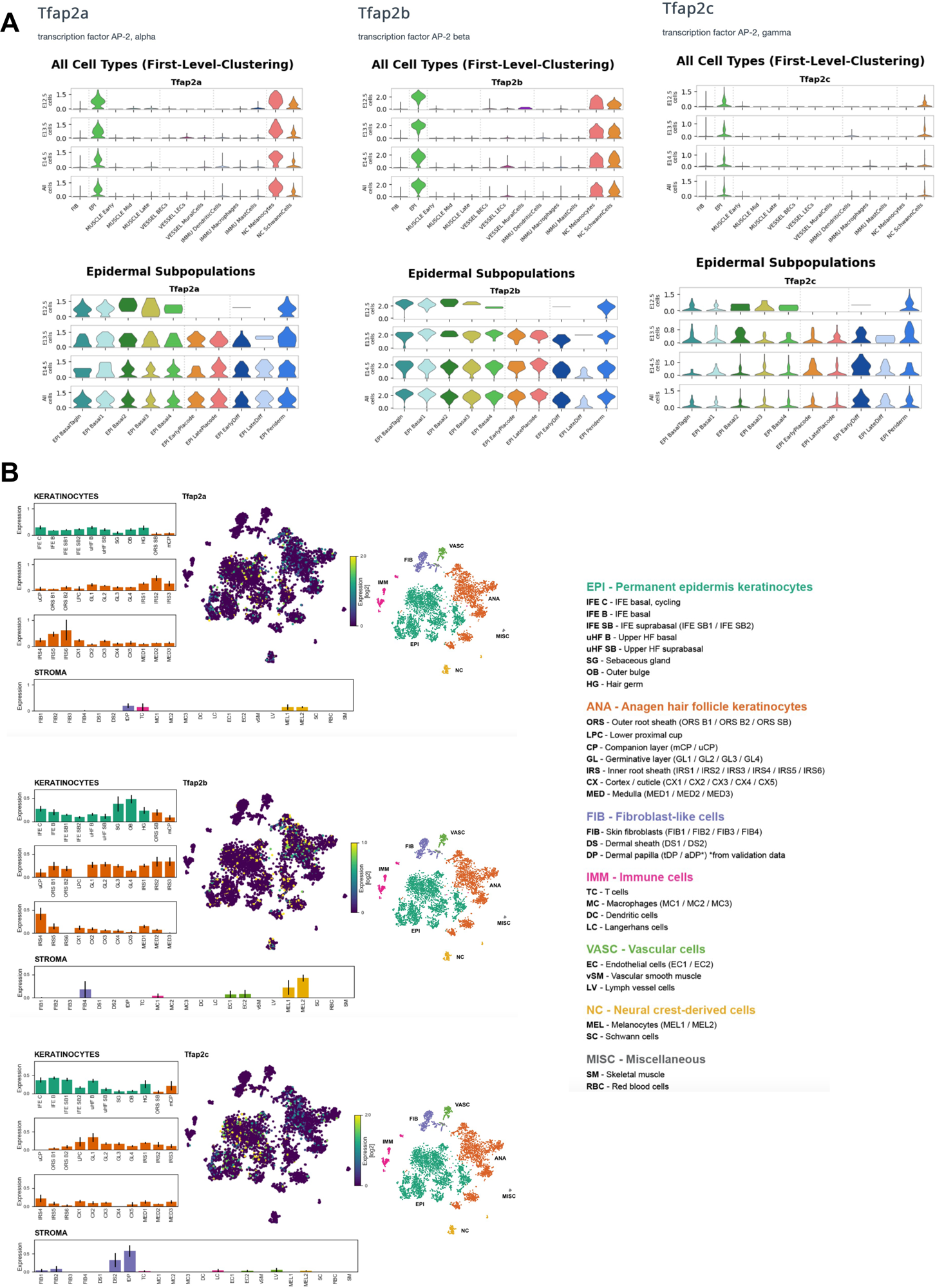
Expression of *Tfap2a*, *Tfap2b*, and *Tfap2c* in developing and adult mouse skin. A. Expression of *Tfap2a*, *Tfap2b*, and *Tfap2c* in the developing mouse skin based on scRNA-seq data (from http://kasperlab.org/embryonicskin; (Jacob et al., 2023)). B. Expression of *Tfap2a*, *Tfap2b*, and *Tfap2c* in the adult mouse skin based on scRNA-seq data (from (Joost et al., 2020)).

**Figure S2:**
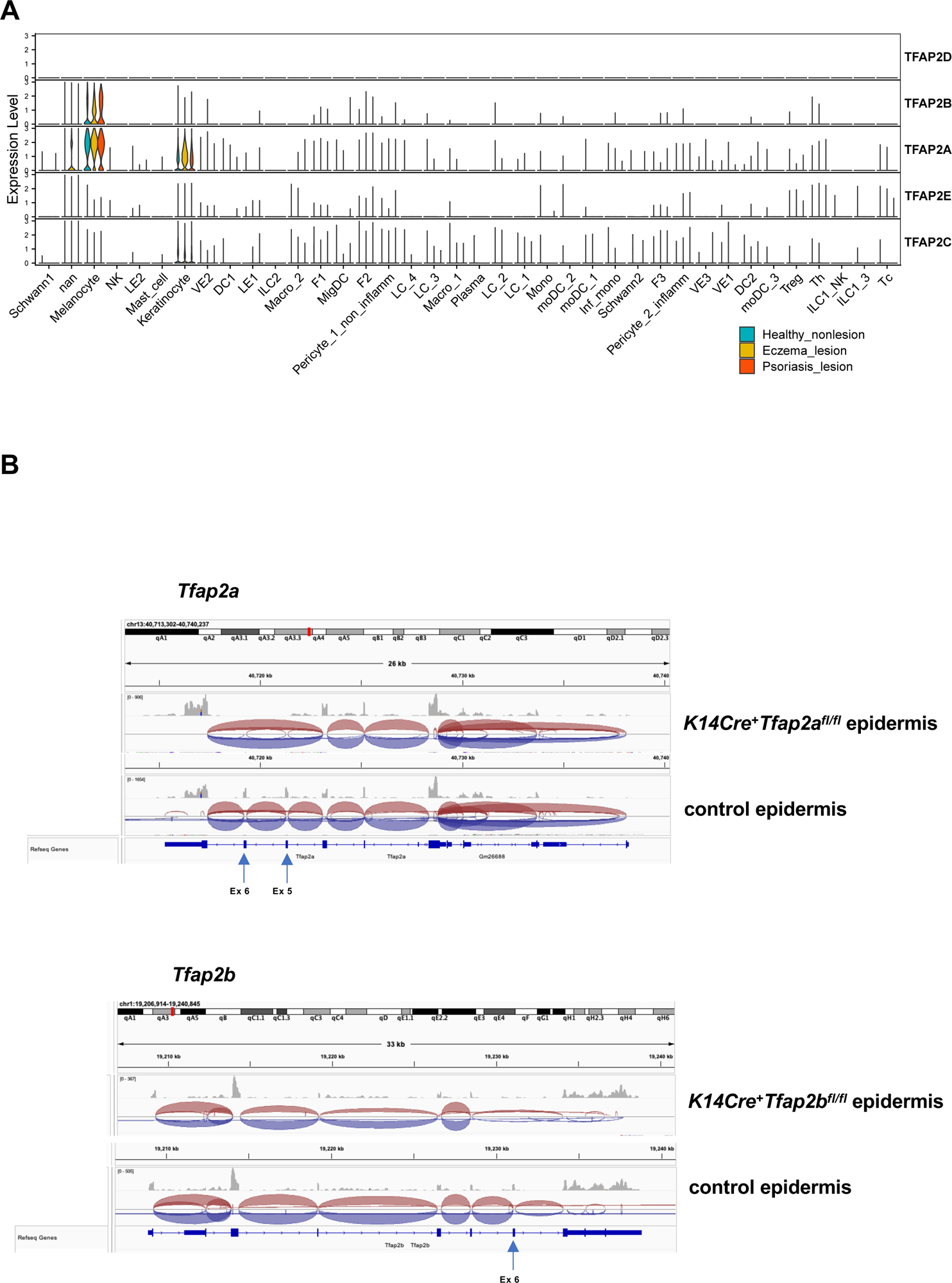
Targeting AP–2α and AP–2β in keratinocytes. A. Violin plots show expression of AP-2 transcription factors in healthy adult human skin based on scRNA-seq data (Reynolds et al., 2021). F, fibroblast; VE, vascular endothelium; LE, lymphatic endothelium; ILC, innate lymphoid cell; NK, natural killer cell; Tc, cytotoxic T cell; T_H_, T helper cell; T_reg_, regulatory T cell; Mac, macrophage; Inf., inflammatory; DC, dendritic cell; LC, Langerhans cell; Mono mac, monocyte-derived macrophage; Mig., migratory; MoDC, monocyte-derived dendritic cell; nan: undefined cell type. B. RNA-seq data from 3-week-old back skin epidermis shows efficient K14Cre-mediated excision of floxed exons 5 and 6 of the *Tfap2a* gene in *K14Cre^+^Tfap2a^fl/fl^*mice and of exon 6 of the *Tfap2b* gene in *K14Cre^+^Tfap2b^fl/fl^*mice.

**Figure S3:**
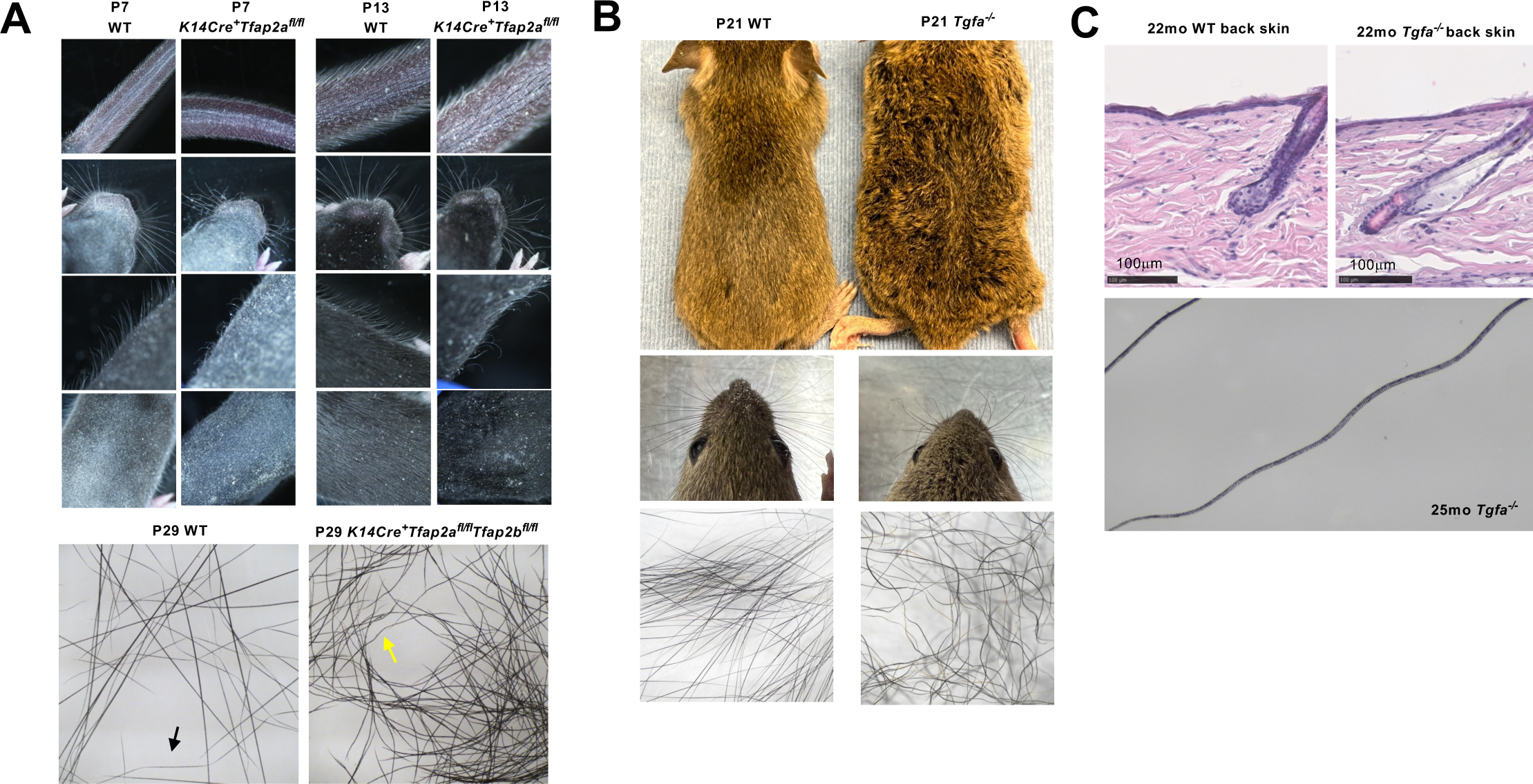
Hair defects due to AP–2α loss are distinct from those due to Tgfa loss. A. P7 and P13 *K14Cre^+^Tfap2a^fl/fl^* mice and control littermates are shown. AP–2α inactivation in keratinocytes results in delayed hair growth and abnormal morphology of hairs and whiskers. Hairs show irregular morphology and loss of zigzag hairs in P29 *K14Cre^+^Tfap2a^fl/fl^Tfap2b^fl/fl^* mice (yellow arrow). Typical zigzag hairs seen in WT controls (black arrow). B. *Tgfa^-/-^* mice show a wavy fur and irregularly curved whiskers, but the hair morphology is distinct from hairs of *K14Cre^+^Tfap2a^fl/fl^*mice. C. No acanthosis or inflammation is observed in the skin of *Tgfa^-/-^* mice (H&E images of 22-month-old back skin). Wavy hairs are maintained in *Tgfa^-/-^* mice also with advanced age. Scale bars, 100 mm.

**Figure S4:**
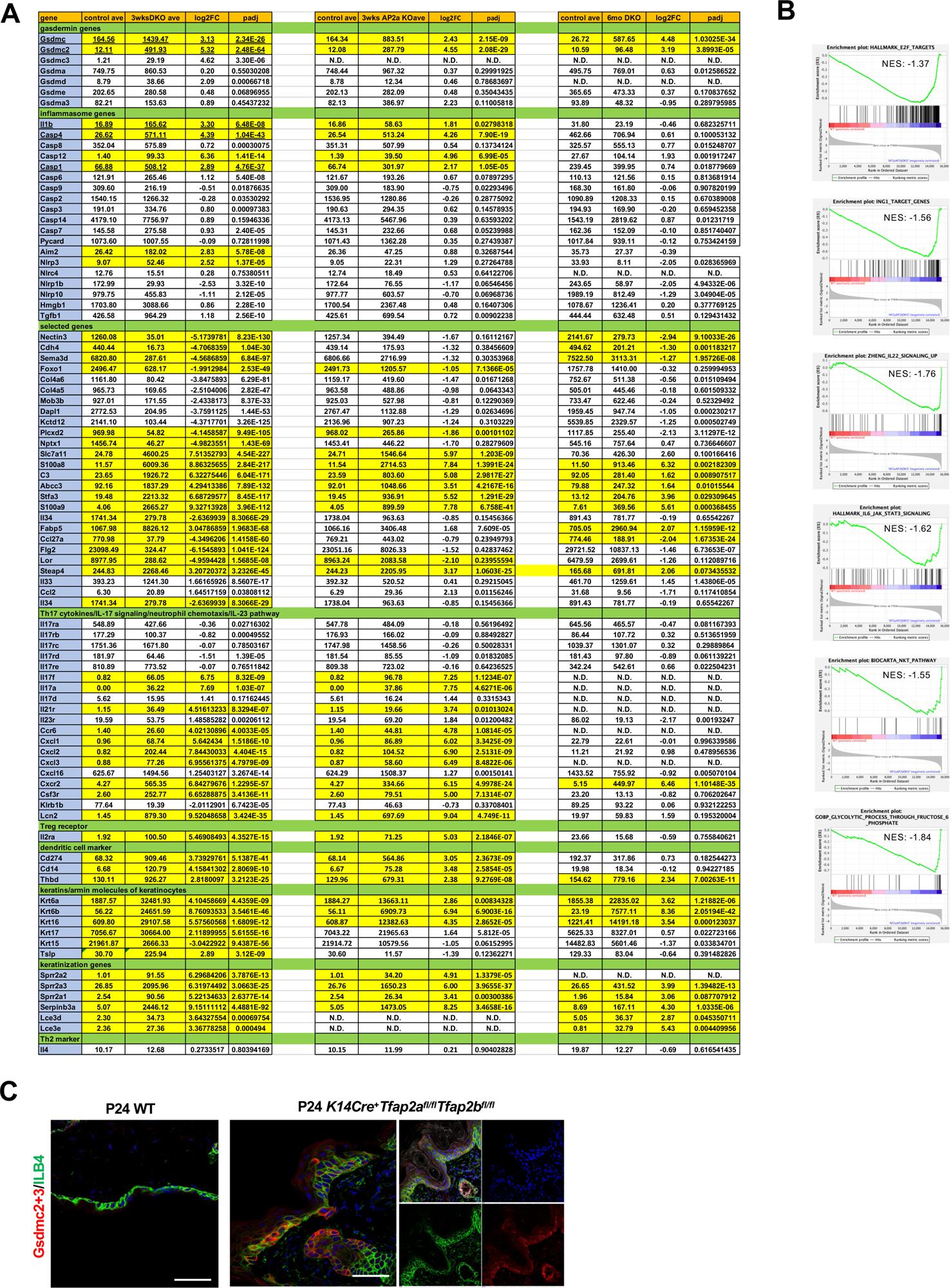
Comparison of key differentially expressed genes between transcriptomes of epidermis lacking AP–2α or AP–2α/AP–2β. A. Selected genes shown from RNA-seq data of back skin epidermis of 3-week-old *K14Cre^+^Tfap2a^fl/fl^Tfap2b^fl/fl^*mice, *K14Cre^+^Tfap2a^fl/fl^* mice, as well as 6-month-old *K14Cre^+^Tfap2a^fl/fl^Tfap2b^fl/fl^* mice (relative to their age-matched WT controls). Transcript levels, log2FC, and padj are shown. B. GSEA of RNA-seq data from back skin epidermis of 3-week-old *K14Cre^+^Tfap2a^fl/fl^Tfap2b^fl/fl^*mice compared to WT controls. Enrichment plots of selected pathways are shown. NES: normalized enrichment scores. Additional results shown in Figure 5B. C. Immunolabeling shows increased Gsdmc2/3 (red) in keratinocytes of *K14Cre^+^Tfap2a^fl/fl^Tfap2b^fl/fl^* mice or *K14Cre^+^Tfap2a^fl/fl^* mice. Scale bars: 50 mm.

**Figure S5:**
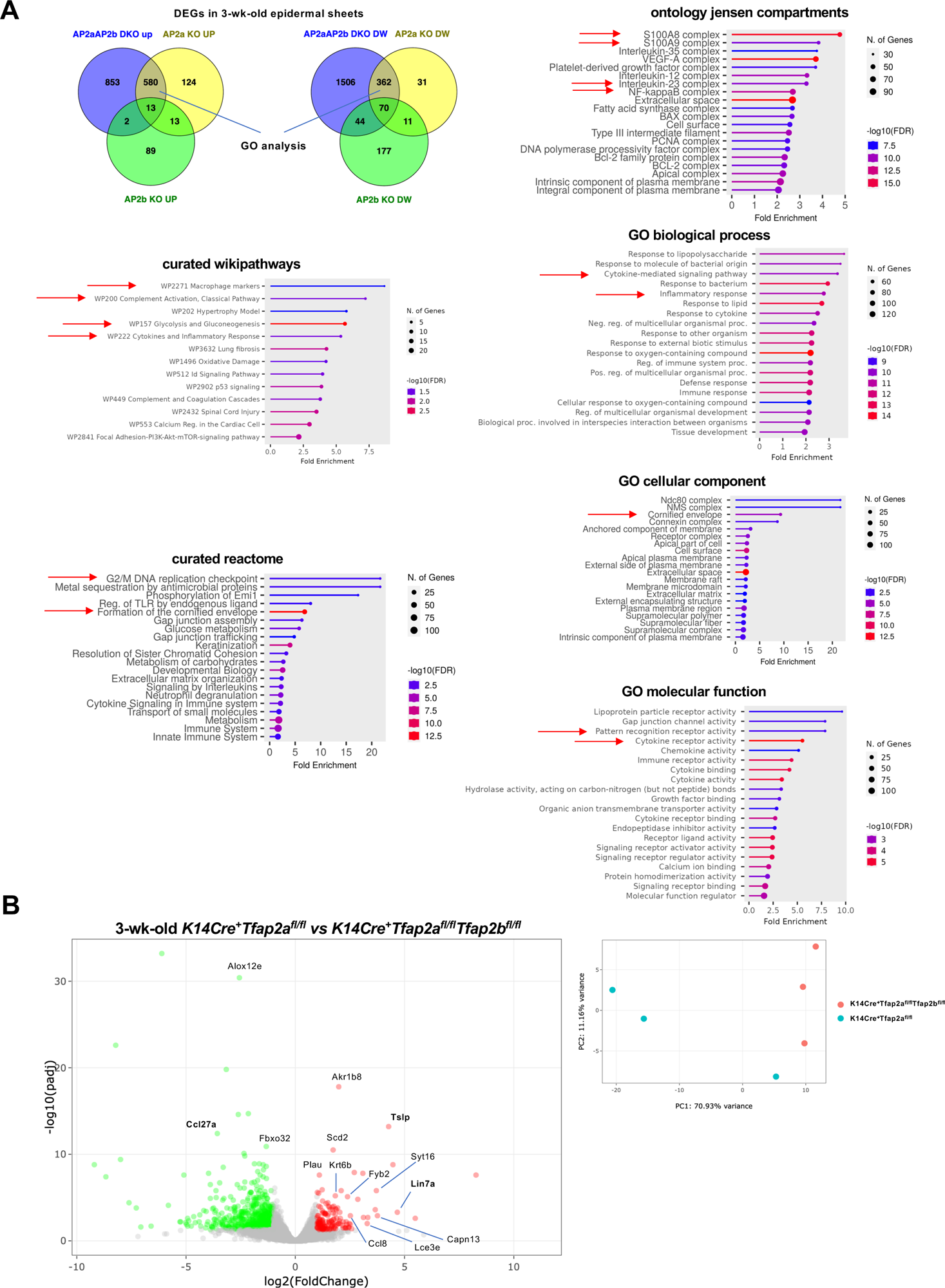
Loss of AP–2α in keratinocytes results in an activation of multiple inflammatory pathways. A. Functional pathway analyses of DEGs that are up- or downregulated in the epidermis of both 3-week-old *K14Cre^+^Tfap2a^fl/fl^Tfap2b^fl/fl^*mice and *K14Cre^+^Tfap2a^fl/fl^* mice. Arrows indicate increased activation of inflammatory pathways, including IL-17 signaling, and other selected key pathways. B. Left: Volcano plot of RNA-seq data show DEGs expressed in back skin epidermis of 3-week-old *K14Cre^+^Tfap2a^fl/fl^* mice compared to *K14Cre^+^Tfap2a^fl/fl^Tfap2b^fl/fl^* mice. Significantly upregulated DEGs in double mutant mice in red and downregulated DEGs in green. Selected genes are shown. Right: PC analysis of these RNA-seq data.

**Figure S6:**
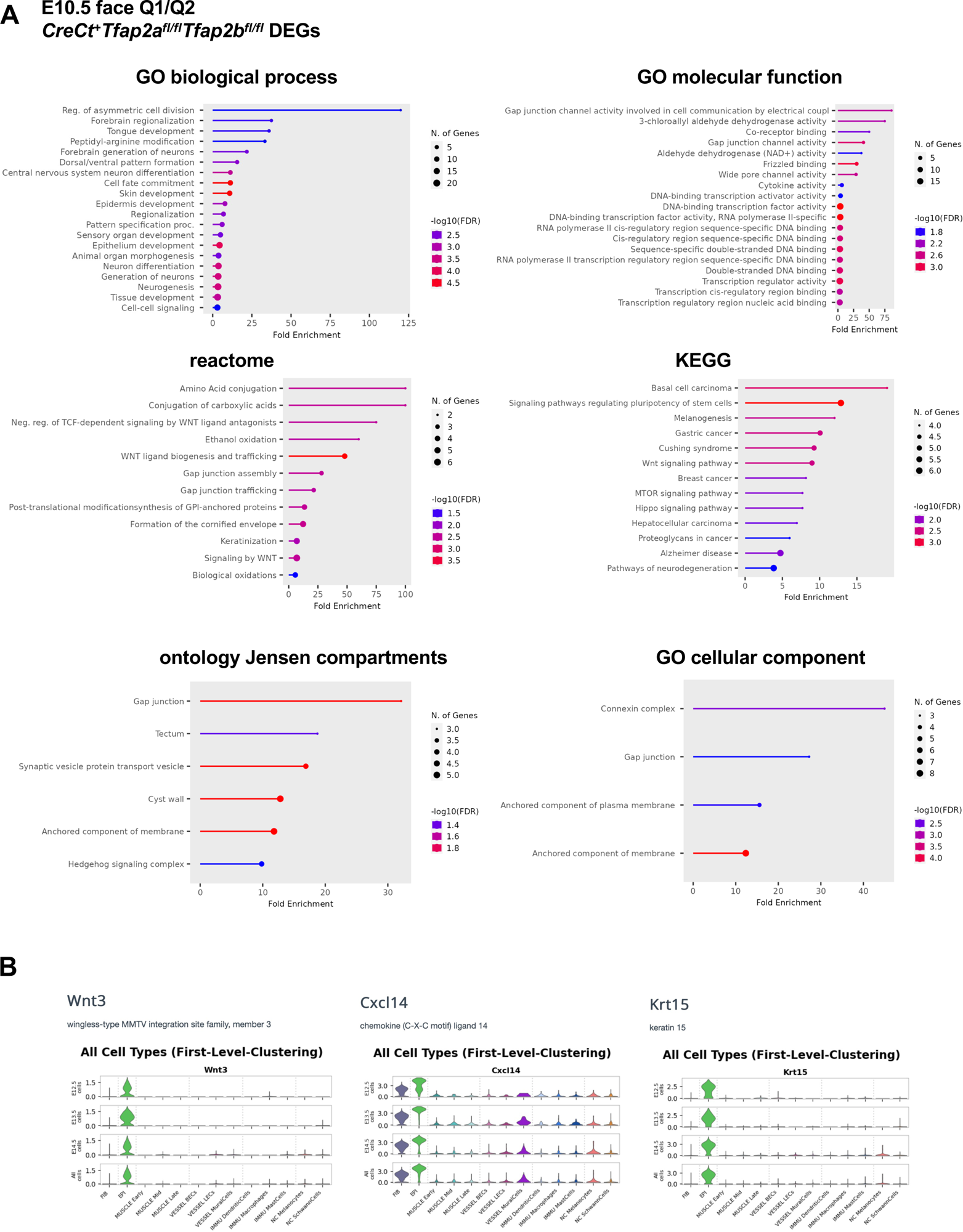
GO analysis of DEGs in E10.5 embryonic mouse heads in which AP–2α and AP–2β were inactivated at ∼E8.5 in surface ectoderm. A. Crect^+^*Tfap2a^fl/fl^Tfap2b^fl/fl^* mice with a focus on epidermis-derived genes (Q1/Q2) in Crect^+^*Tfap2a^fl/fl^Tfap2b^fl/fl^*mice, as described in (Van Otterloo et al., 2022). B. *Wnt3*, *Cxcl14*, and *Krt15* are surface ectoderm markers expressed early during mouse epidermis development. Expression of *Wnt3*, *Cxcl14*, and *Krt15* in the developing mouse skin based on scRNA-seq data (from http://kasperlab.org/embryonicskin; (Jacob et al., 2023)).

**Figure S7:**
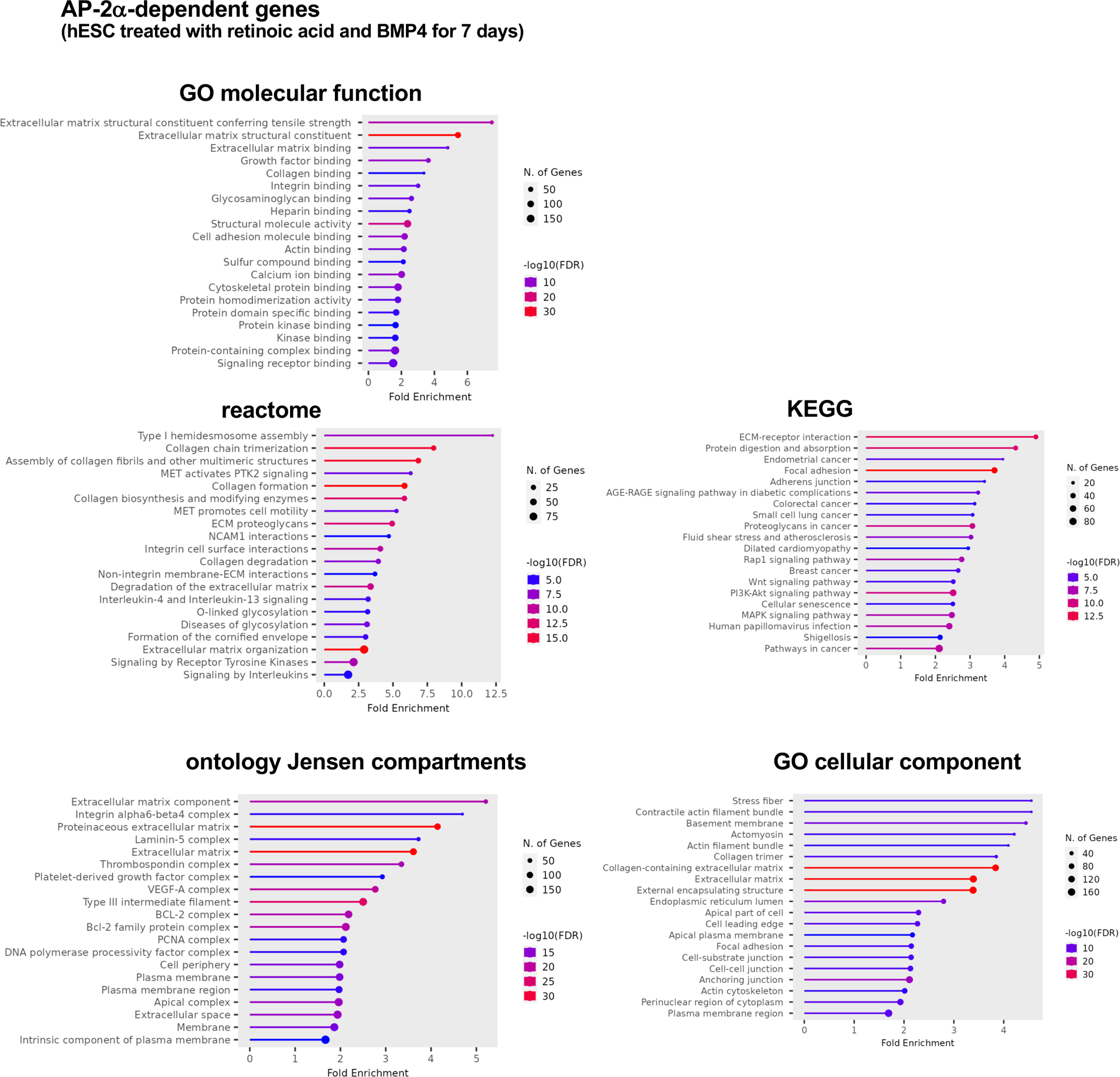
GO analysis of DEGs associated with AP–2α-dependent genes in surface ectoderm cells derived from hESC. Human ES cells (hESC) treated with retinoic acid and BMP4 for 7 days, as described (Collier et al., 2023).

**Figure S8:**
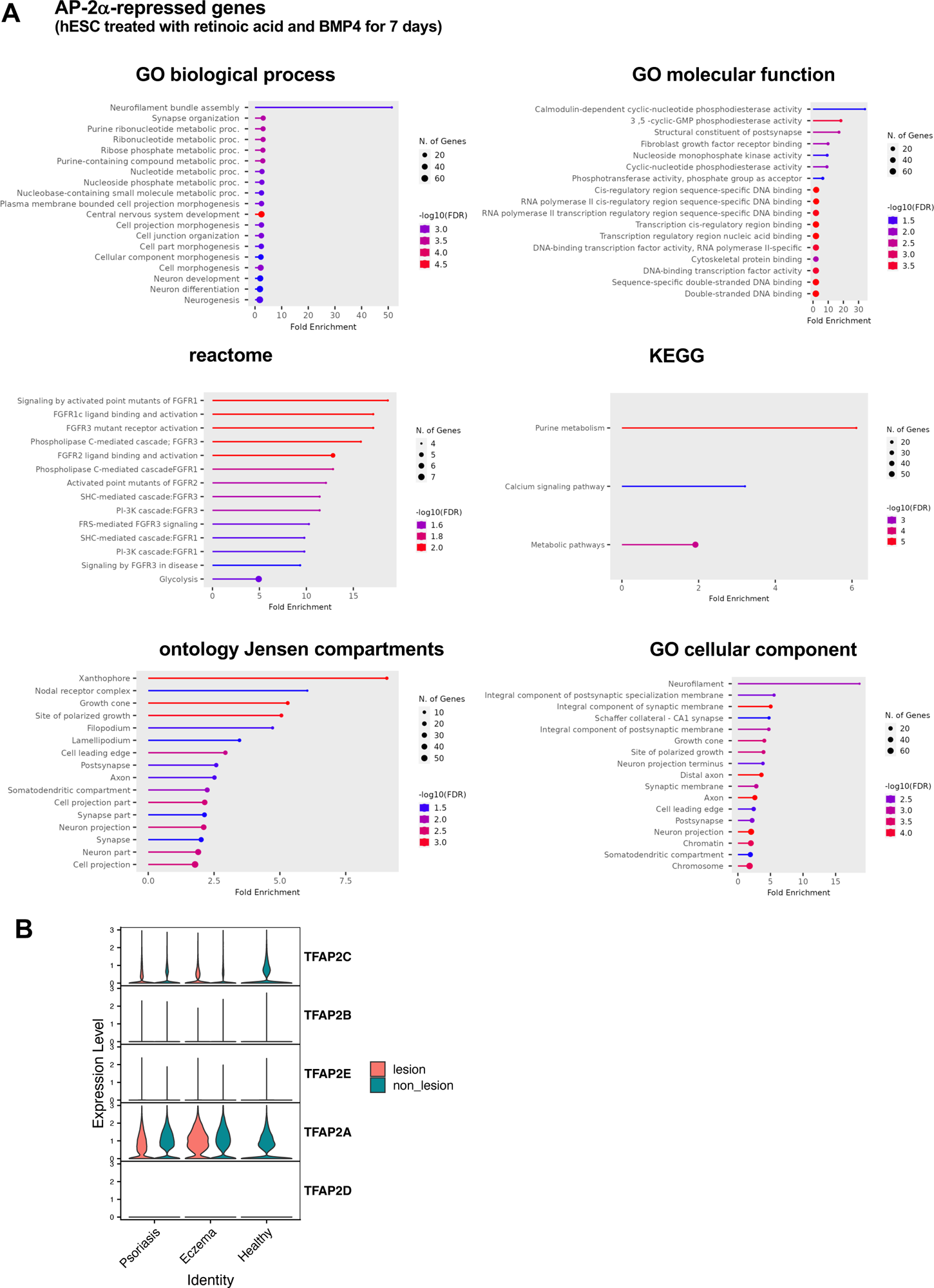
GO analysis of DEGs associated with AP–2α-repressed genes in surface ectoderm cells derived from hESC. A. GO analysis of DEGs associated with AP–2α-repressed genes in surface ectoderm cells derived from hESC treated with retinoic acid and BMP4 for 7 days, as described (Collier et al., 2023). B. AP-2 transcription factor expression in keratinocytes from healthy human skin and from lesional and non-lesional psoriasis or eczema (atopic dermatitis) skin. Derived from scRNA-seq data (Reynolds et al., 2021).

**Table S1:**

**RNA-seq data of experimental mouse groups.**

**Table S2:**

**Genes representing Venn diagrams in Figures 7G and 7H.**

**Comparison groups:**

1. AP2a-dep or AP–2αrep.: DEGs associated with AP–2α-repressed or AP–2α-dependent genes in surface ectoderm cells derived from hESC treated with retinoic acid and BMP4 for 7 days, as described (Collier et al., 2023).
2. E10: E10.5 face from Crect^+^*Tfap2a^fl/fl^Tfap2b^fl/fl^* mice with a focus on epidermis-derived genes (Q1/Q2) in Crect^+^*Tfap2a^fl/fl^Tfap2b^fl/fl^*mice, as described in (Van Otterloo et al., 2022).
3. DKO: DEGs from epidermis 3-week-old *K14Cre^+^Tfap2a^fl/fl^Tfap2b^fl/fl^* mice.
4. AD: ScRNA-seq data of keratinocytes from healthy keratinocytes from lesional atopic dermatitis (eczema) skin (Reynolds et al., 2021).
5. PSX: ScRNA-seq data of keratinocytes from healthy keratinocytes from lesional psoriasis skin (Reynolds et al., 2021).

